# Remyelination restores myelin content on distinct neuronal subtypes in the cerebral cortex

**DOI:** 10.1101/2021.02.17.431685

**Authors:** Cody L. Call, Dwight E. Bergles

## Abstract

Axons in the cerebral cortex exhibit diverse patterns of myelination, with some axons devoid of myelin, some exhibiting discontinuous patches of myelin, and others continuous myelin that is interrupted only by nodes of Ranvier. Oligodendrocytes establish this pattern by sorting through a high density of potential targets to select a small cohort of axons for myelination; however, the myelination patterns established on distinct excitatory and inhibitory neurons within the cortex remain to be fully defined and little is known about the extent to which these patterns are restored after oligodendrocyte regeneration. Here we show that axons in layer I of the somatosensory cortex, a key region for integration of input from local and distant sources, exhibit an extraordinarily diverse range of myelination patterns, even among distinct neuronal subtypes. Although larger axons were more often selected for myelination, neuronal identity profoundly influenced the probability of myelination. The relative differences in myelination among neuron subtypes were preserved between cortical areas with widely varying myelin density, suggesting that regional differences in myelin abundance arises through local control of oligodendrogenesis, rather than selective reduction of myelin on distinct neuron subtypes. By following the loss and regeneration of myelin sheaths along defined neurons *in vivo* we show that even though the distribution of myelin on individual PV and VM neuron axons was altered following remyelination, the overall myelin content on these neurons was restored. The findings suggest that local changes in myelin can be tolerated, allowing opportunistic selection of available targets by newly formed oligodendrocytes to restore relative differences in myelin content between functionally distinct neurons.

## INTRODUCTION

Circuits within the cerebral cortex are formed by a diverse population of neurons that integrate input from both local axon collaterals and long-range projections within specific lamina. In the upper most layer of the cortex (layer I), axonal projections synapse onto the dendritic tufts of pyramidal neurons, serving as a critical hub for integrating local inhibition, thalamocortical and corticocortical excitation, and long-range neuromodulation, providing top-down regulation of sensory inputs^1–3^. Despite the critical need to integrate input from both local and distant inputs in this region, and the critical ability of myelin to control timing of synaptic activity, we have only a limited understanding of the myelin patterns along axons of distinct neuron subtypes that course within this cortical area. Although a small fraction of these axons are well-myelinated and extend within layer I for many millimeters in rodents (or centimeters, in primates) before terminating^4,5^, the overall myelin content within each cortical region is small^6^, suggesting a high level of specificity in oligodendrocyte selection of axons. Approximately 10% of myelin sheaths in layer I surround inhibitory axons, the majority of which express parvalbumin (PV)^7,8^, but the vast majority are associated with an unknown population of neurons. Moreover, individual axons in the cortex can vary extensively in the number and pattern of myelin sheaths along their lengths^9–11^. This “discontinuous” myelination has been well-described on PV interneurons and proximal axon segments of pyramidal neurons^8,11^, but the pattern of myelin along these and other types of neurons has not been explored in layer I.

Schwann cells in the peripheral nervous system select axons for myelination strictly by axon diameter^12^, with a threshold of ∼1.5 μm^12–14^. Similarly, oligodendrocytes in the CNS only myelinate axons or inert fibers *in vitro* with diameters > 0.3 μm^15–17^, suggesting that there are strict biophysical constraints on which axons can be wrapped. Nevertheless, individual oligodendrocytes are capable of myelinating a range of axon sizes^18^ and provide sheaths of different thicknesses and lengths, often correlated with the diameter of each axon^18,19^, illustrating the additional complexity of CNS myelination. The discontinuity of myelin along individual axons and the numerous axons of permissive diameter that lack myelin indicate that oligodendrocytes in gray matter may be influenced by other factors in addition to axon diameter, such as spiking activity^20–24^ or surface protein expression^25^ to yield cell type specific myelination patterns.

The complex patterns of myelination that exist in the cortex create significant challenges for repair. Demyelinating lesions of the gray matter in multiple sclerosis (MS) most commonly occur just below the pia^26–28^, and are associated with cognitive disabilities and poorer prognosis^26,29–33^, making this region of particular interest for understanding the identities of the axons myelinated and the extent to which they are remyelinated in recovery. Cortical PV interneurons become extensively myelinated very early in development^34^, which is thought to be important for preventing runaway excitation in nascent circuits and for coordinating pyramidal neuron firing to generate high frequency oscillations^35,36^. Recent *in vivo* imaging studies indicate that the distribution of oligodendrocytes and the overall pattern of myelin changes upon recovery from cuprizone-induced demyelination^37^, but it is not known if this reorganization affects the distribution and content of myelin along excitatory and inhibitory neurons equally. Myelin within layer I of the somatosensory cortex is completely restored after cuprizone induced destruction of oligodendrocytes^37^, providing a unique opportunity to study the specificity of the remyelination process *in vivo*.

Here we used a combination of cell specific axon labeling with high resolution imaging of myelin sheaths to define the myelination patterns of seven distinct excitatory and inhibitory neuronal subtypes that extend axon collaterals within layer I of the somatosensory cortex. This analysis revealed a diversity of myelin patterns that did not closely follow functional class (excitatory versus inhibitory) or cell body location (thalamus versus cortex), with discontinuous myelination observed on all myelinated axons in these adult mice. The probability of myelination could be predicted through a combination of cell type and axon diameter, indicating that the observed patterns are strongly influenced by cell intrinsic factors rather than just cell morphology. By performing longitudinal time lapse imaging of local PV interneurons and thalamocortical VM neurons we found that although the precise pattern of myelin along individual axons was altered following regeneration, at the population level, the total myelin content on these distinct neuronal subtypes was preserved. These findings suggest that regeneration of oligodendrocytes relies on opportunistic target selection to restore the content of myelin on diverse neuronal subtypes in the mammalian cortex.

## RESULTS

### Cortical axons exhibit diverse myelination patterns

To define the pattern of myelination along axons of different subtypes of neurons in the cerebral cortex, we used Cre-lox and viral expression strategies to fluorescently label seven neuronal subpopulations that extend axon collaterals into layer I of primary somatosensory cortex (S1) (**Fig. 1a, Table 1**). We compared myelin along axons of two types of GABAergic neurons (parvalbumin (PV), somatostatin (SOM)) and three corticocortical projecting neurons (layer VI corticothalamic: *NTSR1-Cre*^38–40^; layer VIb subplate: *NXPH4-Cre*^41,42^; layer Va/Vb pyramidal: *RBP4-Cre*^43^) by crossing to fluorescent reporter mice (*Ai3, Ai9*)^44^. These Cre lines have been well-characterized in previous studies and are highly selective in labeling pyramidal neurons within their corresponding lamina^45,46^ (**Extended Data Fig. 1a-h**). In addition, two thalamocortical projections, the ventral medial nucleus (VM) and medial posterior nucleus (PO, were analyzed by injecting tdTomato expressing AAV virus within different regions of the thalamus. Horizontal sections that preserved the orientation of axonal trajectories (parallel to the pia) were then collected from these mice and immunostained for MBP (**Fig. 1b,c**). The complete morphologies of randomly-selected axons were traced within a 675 μm x 675 μm x 40 μm volume and the percent of axon length myelinated (PLM) calculated ([total length of all MBP+ internodes / total axon length] x 100%) (**Fig. 1d–f, Supplementary Video 1**). PLM distributions varied considerably between the six neuronal subtypes that were myelinated (**Fig. 2a**); NTSR1 expressing pyramidal neurons were not myelinated. Moreover, axons of each neuronal subtype ranged from unmyelinated to nearly completely myelinated, revealing the high variability in cortical myelination even within the same neuronal subtype. Although differences in the degree of axonal collateralization between neuronal subtypes influenced the total axon length measured (**Fig. 2b**), this variability did not correlate with PLM for any neuronal subtype except SOM (R^2^ = 0.373) (**Extended Data Fig. 1**), suggesting that under sampling did not strongly influence detection of these myelination patterns.

**Figure 1.**
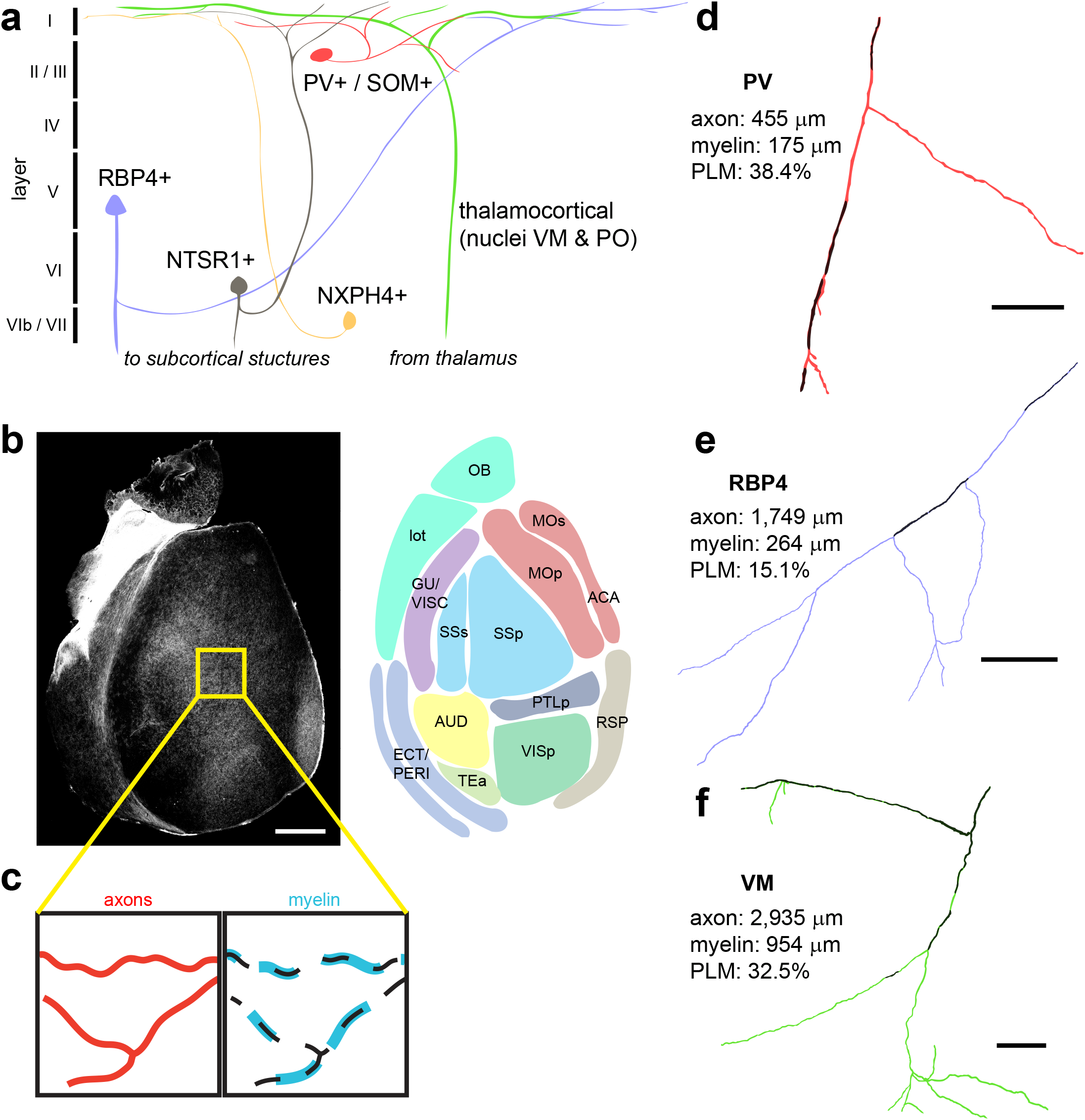
High resolution tracing of distinct axon populations reveals discontinuous myelination patterns. **a**, Schematic of cortex indicating the sources of axons from the distinct neuronal populations investigated. Note that the axons depicted may not necessarily be the primary projection of each neuron. **b**, A horizontal section of the entire cortical flatmount and schematic representing approximate cortical areas. **c**, Schematic of experimental design. Traces (black dotted line) of fluorescently-labeled axons (red) were used to trace corresponding MBP-immunostained myelin sheaths (cyan). **d–f**, Full reconstructions of traced PV (**d**), RBP4 (**e**), and VM (**f**) axon morphologies within cortical layer I and their associated dimensions (PLM, percent length myelinated). Black myelin sheaths are overlaid on colored axons. Scale bars, 100 μm. Abbreviations: OB, olfactory bulb; lot, lateral olfactory tract; MOp/MOs, primary/secondary motor; ACA, anterior cingulate area; GU/VISC, gustatory/visceral; SSp/SSs, primary/secondary somatosensory; PTLp, posterior parietal association; AUD, auditory; RSP, retrosplenial; VISp, primary visual; ECT/PERI, ectorhinal/perirhinal; TEa, temporal association.

**Table 1.**
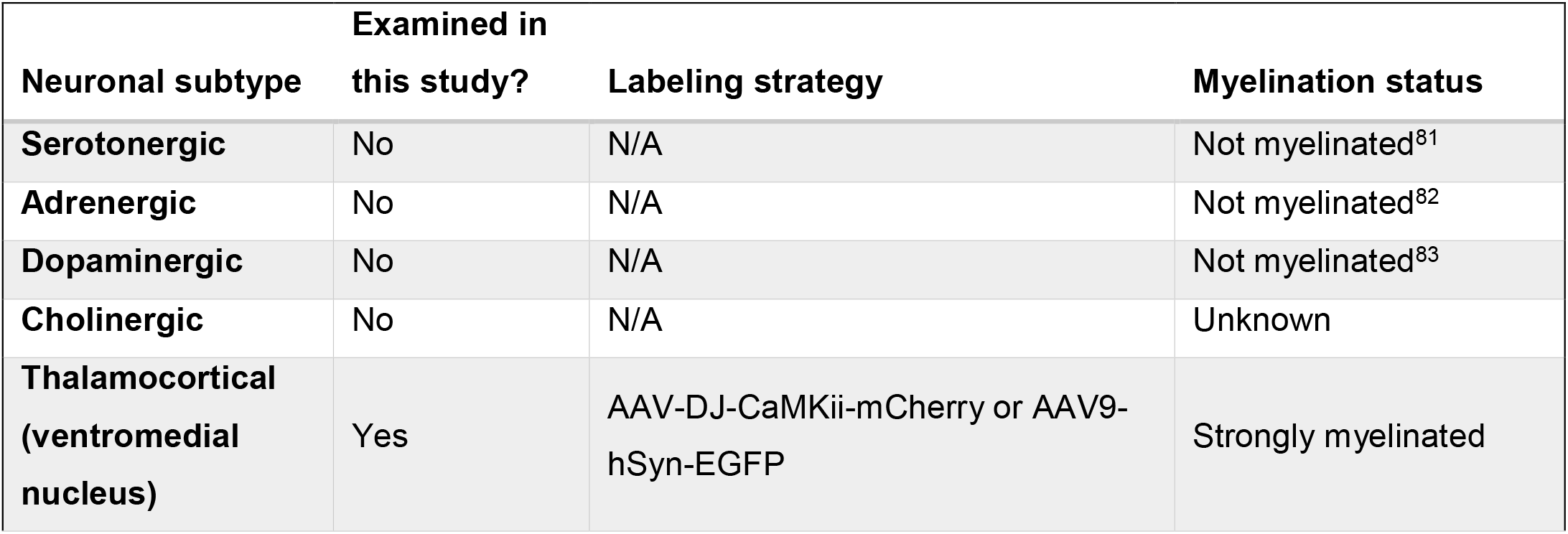
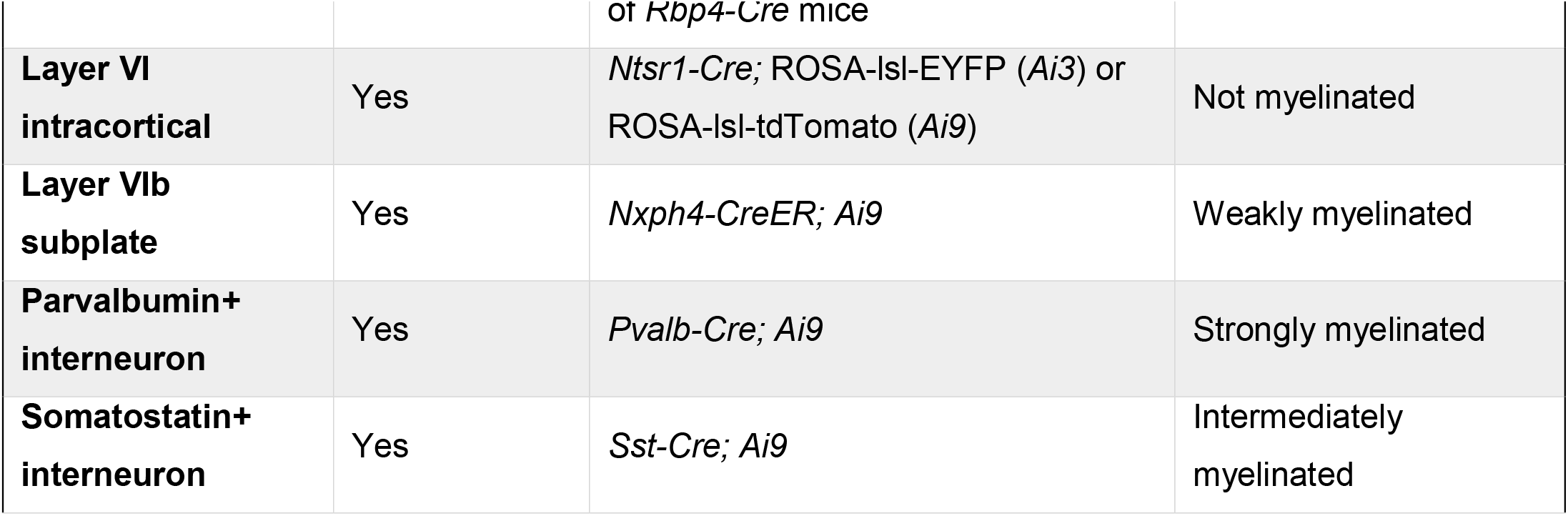
Layer I-projecting neuronal subtypes and labeling strategies

**Figure 2.**
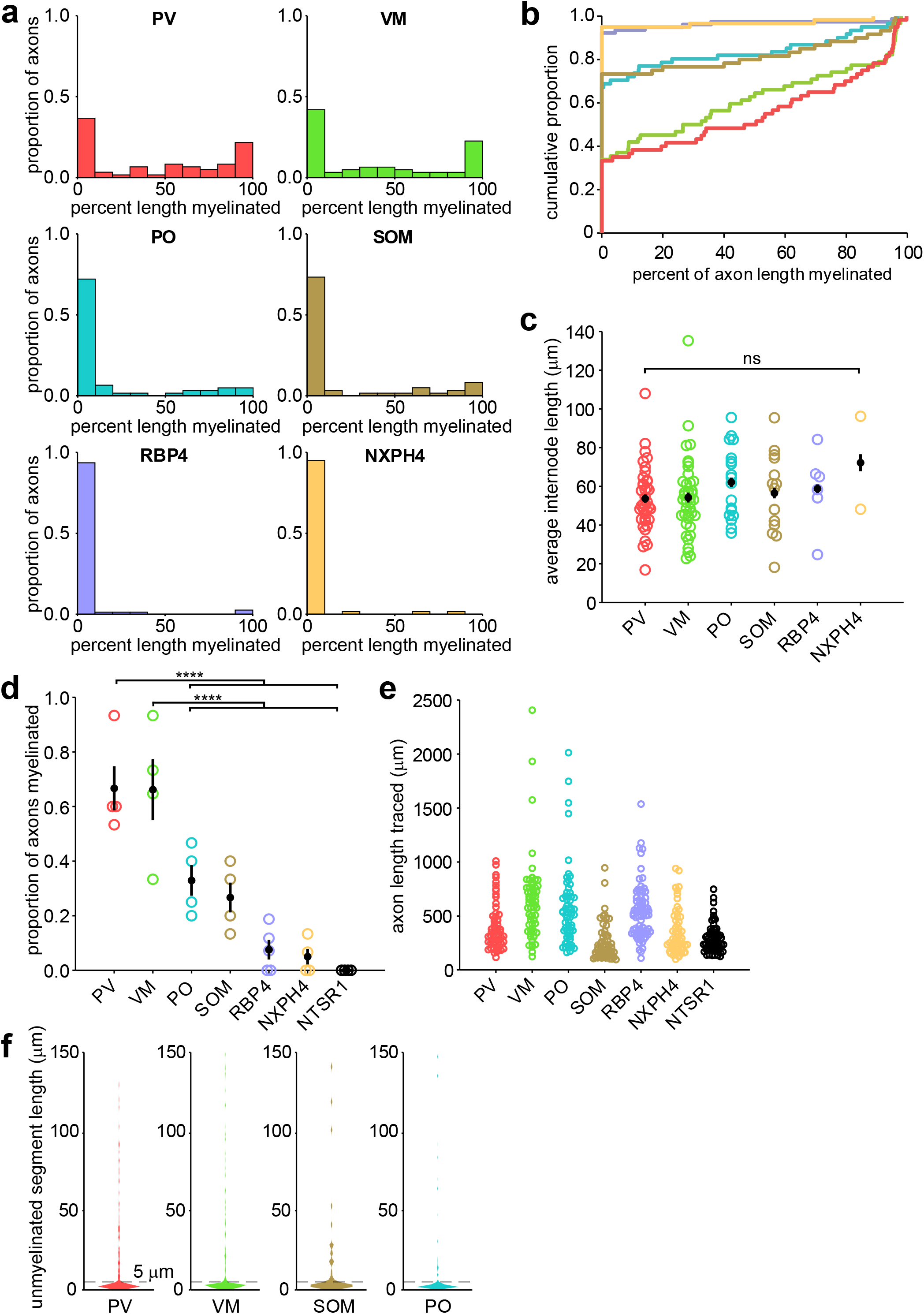
Different neuronal subtypes exhibit diverse myelination profiles. **a**, Myelination profiles for the six neuronal subtypes examined in this study with myelinated axons. Each plot is a histogram of pooled data across four animals for each subtype representing the proportion of axons with different percent length myelinated in 10% bins (n = 60 (PV), 62 (VM), 60 (SOM), 61 (PO), 60 (NXPH4), 78 (RBP4), 60 (NTSR1) axons). See **Table 2** for statistical comparisons. **b**, Cumulative distribution plot of data in (**a**), showing three distinct classes of myelination patterns. **c**, Average internode lengths per axon for each neuronal subtype. **d**, Proportions of axons myelinated per neuronal subtype (N = 4 mice for each subtype). **e**, Lengths for each axon traced (same Ns as in (**a**)). **f**, Violin plots of the lengths of unmyelinated axon segments (axonal distance between two consecutive MBP segments) for the top four most strongly myelinated neuronal subtypes. Unmyelinated segments of < 5 μm (typical maximum length of nodes of Ranvier) are most common, while unmyelinated segments between 5 and 150 μm are uniformly distributed.

As previously described^7,8^, PV axons were among the most highly myelinated, with approximately two thirds of traced axons associated with at least one internode (average proportion: 0.7 ± 0.08, n = 4 mice) and about half of PV axons exhibiting a PLM > 50% (**Fig. 2a,b**). VM thalamocortical axons were similarly well-myelinated (average proportion: 0.6 ± 0.1, n = 4 mice) and exhibited similar myelination patterns as PV axons (*p* = 0.99, Kruskal-Wallis one-way ANOVA with Dunn–Šidák correction for multiple comparisons) (**Fig. 2a, Table 2**). All other axons examined were myelinated much less frequently (average proportions: PO, 0.3 ± 0.06; SOM, 0.3 ± 0.05; RBP4, 0.08 ± 0.04; NXPH4, 0.05 ± 0.03; NTSR1, 0; n = 4 mice each) (**Fig. 2b, Table 2**). There was no difference in the average length of internodes between neuronal subtypes (*p* = 0.514, one-way ANOVA; **Fig. 2c**), indicating that PLM differences reflect the number of internodes per axon.

**Table 2:** Multiple comparison tests for myelination patterns

| Sample 1 | Sample 2 | Lower 95% CI | Estimate | Upper 95% CI | Corrected *p* value |
| --- | --- | --- | --- | --- | --- |
| PV | VM | -42.98 | 5.88 | 54.74 | 1 |
| PV | SOM | 29.57 | 78.83 | 128.09 | **4.19 × 10<sup>-5</sup>** |
| PV | PO | 23.95 | 73.01 | 122.06 | **0.0002** |
| PV | RBP4 | 71.34 | 117.67 | 164.00 | **1.56 × 10<sup>-12</sup>** |
| PV | NXPH4 | 72.66 | 121.92 | 171.18 | **6.43 × 10<sup>-12</sup>** |
| VM | SOM | 24.09 | 72.95 | 121.81 | **0.0002** |
| VM | PO | 18.47 | 67.12 | 115.78 | **0.0008** |
| VM | RBP4 | 65.88 | 111.79 | 157.69 | **1.51 × 10<sup>-11</sup>** |
| VM | NXPH4 | 67.17 | 116.03 | 164.90 | **5.37 × 10<sup>-11</sup>** |
| SOM | PO | -54.89 | -5.83 | 43.23 | 1 |
| SOM | RBP4 | -7.49 | 38.84 | 85.17 | 0.1921 |
| SOM | NXPH4 | -6.18 | 43.08 | 92.34 | 0.1457 |
| PO | RBP4 | -1.45 | 44.66 | 90.78 | 0.0664 |
| PO | NXPH4 | -0.15 | 48.91 | 97.97 | **0.0514** |
| RBP4 | NXPH4 | -42.08 | 4.25 | 50.58 | 1 |

Three general myelination profiles were evident in the PLM distributions: strongly myelinated (PV and VM), intermediately myelinated (SOM and PO), and weakly myelinated (RBP4 and NXPH4). Overall, PO and SOM axons shared similar myelination patterns, matching that observed in deeper cortical layers^47^, as did RBP4 and NXPH4 axons. Notably, these myelination classes did not match functional classifications: while all corticocortical axons were very weakly myelinated, the interneuronal and thalamocortical axons were each split between strongly and intermediately myelinated classes.

Myelination along individual cortical axons is often discontinuous, with myelinated segments interrupted by long stretches of unmyelinated axon^9–11^. To determine the extent of myelin discontinuity along different axons, we quantified the lengths of unmyelinated segments among the well-myelinated neuronal subtypes (PV, VM, SOM and PO). To reach the same PLM, axons could be myelinated with frequent gaps between adjacent internodes or have infrequent, longer gaps between local continuously myelinated regions, with the only interruptions arising from nodes of Ranvier. Plots of unmyelinated segment length revealed that all axons exhibited a large cluster of short lengths typical for nodes of Ranvier (0-5 μm; PV: 67%, VM: 73%, PO: 66%, SOM: 69% of all lengths) (**Fig. 2f**), indicating that internodes tend to be clustered along axons to form continuously myelinated segments. Together, these studies reveal that different neurons that extend axons within layer I of the adult somatosensory cortex exhibit distinct, highly variable myelination patterns.

### Axonal myelination patterns scale between cortical regions

The density of myelin varies considerably across the cortical mantle (**Fig. 1b**), with boundaries between cortical areas defined by their distinct myeloarchitecture^48,49^. It is not known if these differences reflect changes in myelin content along certain classes of neurons or proportional decreases in myelin content along all axons. To determine if the diversity in myelin patterns observed in the somatosensory cortex are preserved in other cortical regions, we compared the myelination of PV axons across five cortical areas, as PV neurons exhibit consistent axon density across the primary sensory and motor cortices, with the exception of a slightly lower density in association cortex^50^. Among sensory cortical areas (somatosensory (SS), visual (VIS), and auditory (AUD)), the density of MBP immunoreactivity was similar (mean gray value of binarized z projection, SS: 50 ± 8; VIS: 66 ± 8; AUD: 61 ± 11, arbitrary units) and the PLM distribution of PV axons was not significantly different between these areas (SS vs. VIS: *p* = 0.16; SS vs. AUD: *p* = 0.45; VIS vs. AUD: *p* = 1, Kruskal-Wallis one-way ANOVA with Dunn– Šidák correction for multiple comparisons) (**Fig. 3a–c, Table 3**). As expected, in cortical areas with lower myelin content (i.e. secondary motor (MOs): 42 ± 8; temporal association area (TEa): 14 ± 3), the PLM distribution was significantly left-shifted, with fewer PV axons myelinated (**Fig. 3d,e; Table 3**). However, even in these sparsely myelinated areas, some PV axons were continuously myelinated, suggesting that the preference for this neuronal subtype is maintained across the cortex. Consistent with this hypothesis, when the myelin distribution of PV axons was scaled to overall MBP density (scaled myelination prevalence = average binarized MBP intensity/proportion of axons myelinated) this value was not significantly different between cortical regions (*p* = 0.24, Kruskal-Wallis ANOVA) (**Fig. 3f**). These results suggest that variations in myelin abundance across the cortex reflect regional differences in oligodendrogenesis or oligodendrocyte survival, rather than selective changes in myelin along different classes of neurons.

**Figure 3.**
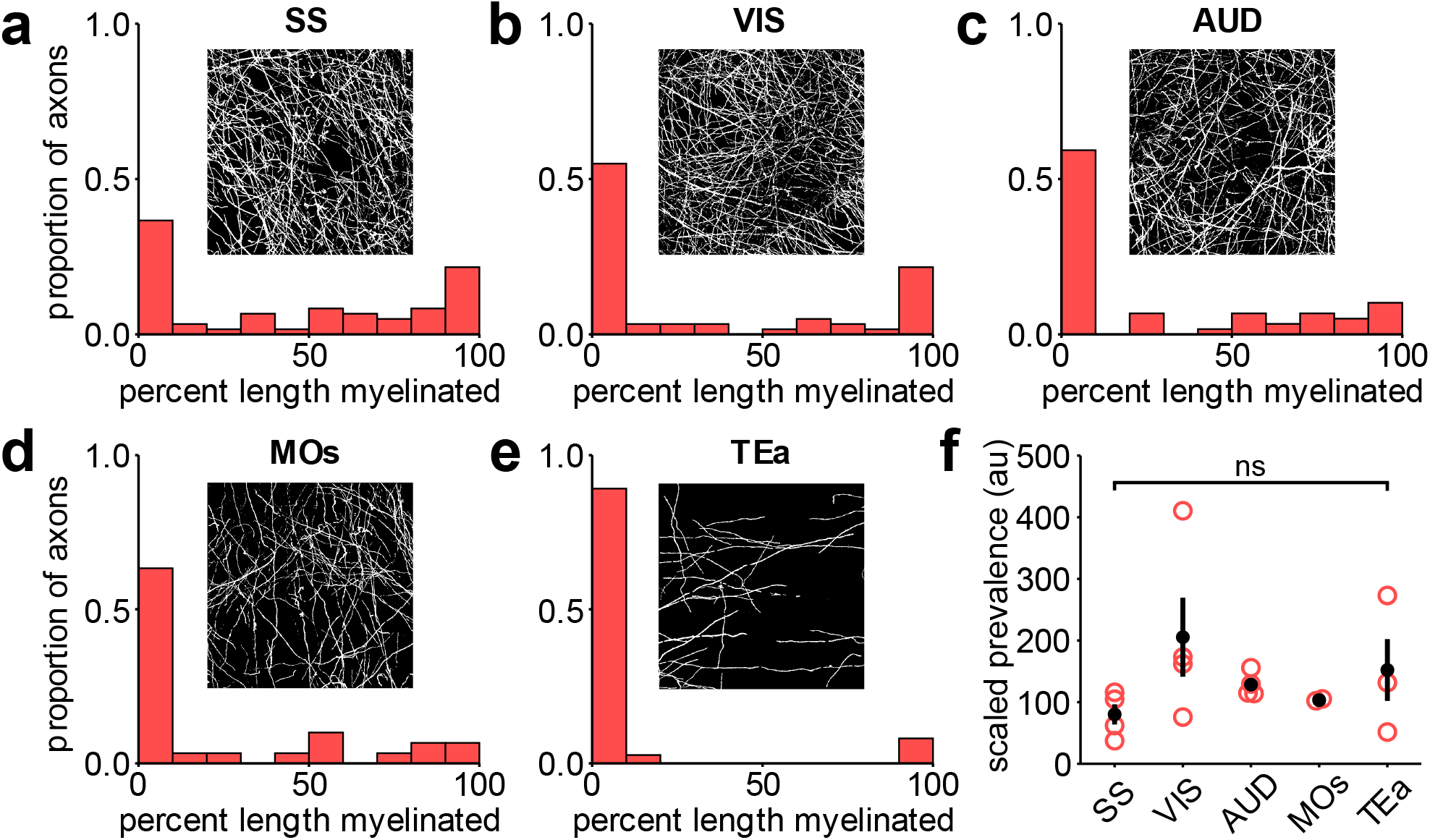
PV axon myelination patterns are consistent across cortical regions. Histogram of percent length myelinated (PLM) for all axons traced in the somatosensory (**a**, N = 60 axons from 4 mice), visual (**b**, N = 60 axons from 4 mice), auditory (**c**, N = 59 axons from 4 mice), secondary motor (**d**, N = 30 axons from 2 mice), temporal association (**e**, N = 37 axons from 3 mice) cortical areas. Insets are representative binarized images of MBP immunostaining from one image stack used for tracing from each area. See **Table 3** for PLM distribution statistics. **f**, Scaled prevalence (MBP intensity ÷ proportion of axons myelinated) of PV axon myelination across all regions analyzed.

**Table 3:** Multiple comparison tests for PV regional myelination

| Sample 1 | Sample 2 | Lower 95% CI | Estimate | Upper 95% CI | Corrected *p* value |
| --- | --- | --- | --- | --- | --- |
| SS | VIS | -5.66 | 32.91 | 71.48 | 0.1567 |
| SS | AUD | -12.33 | 26.08 | 64.49 | 0.4455 |
| SS | MOs | 33.72 | 72.13 | 110.54 | **$1.46 \times 10^{-6}$** |
| SS | TEa | 51.27 | 89.68 | 128.09 | **$6.27 \times 10^{-10}$** |
| VIS | AUD | -45.40 | -6.83 | 31.74 | 1 |
| VIS | MOs | 0.65 | 39.22 | 77.79 | **0.0433** |
| VIS | TEa | 18.20 | 56.77 | 95.34 | **0.0004** |
| AUD | MOs | 7.64 | 46.05 | 84.46 | **0.0079** |
| AUD | TEa | 25.19 | 63.60 | 102.01 | **$3.55 \times 10^{-5}$** |
| MOs | TEa | -20.86 | 17.55 | 55.96 | 0.8937 |

### Axon diameter and neuronal subtype influence myelination patterns

The relationship between axonal diameter and myelination has been thoroughly described in the context of peripheral nerves, CNS white matter, and even primary cell culture^12–15,17,51,52^, establishing that the probability of myelination increases with axonal diameter. To determine if differences in myelination between neuronal subtypes in the cortex are simply due to differences in axon diameter, we quantified the variation in axon diameter among each neuronal subtype in relation to MBP+ myelin sheaths (**Fig. 4a**). Axon dimeters were estimated by measuring the full-width at half maximum (FWHM) of fluorescent intensity (EYFP, tdTomato, or mCherry) across axons using superresolution microscopy (**Fig. 4b**). Across all neuronal subtypes examined there was a positive correlation between axon diameter and probability of being myelinated (*p* = 8.99 × 10^−14^, two-sample Kolmogorov-Smirnov test) (**Fig. 4c,d**). However, for axons with diameters between 0.4 and 0.8 μm (44% of all axons examined) there was remarkable diversity in myelination status among all neuron subtypes, varying from completely myelinated to completely unmyelinated (**Fig. 4c,d**). While the most strongly myelinated subtypes (PV and VM) tended to have larger axon diameters (average axon diameter: PV, 0.6 ± 0.03 μm; VM, 0.5 ± 0.03 μm), their thinnest segments (< 0.5 μm) still had a reasonable likelihood of being myelinated (PV, 30%; VM, 21%) (**Fig. 4c**). To explore relationships between cell identity, axon diameter and myelination status, we fitted a binomial generalized linear model to the entire dataset, whereby the probability of an axon being myelinated is predicted by axon diameter and neuronal subtype (P(myelinated) ∼ 1 + diameter + subtype) (**Table 4**). This analysis revealed that while axon diameter influences myelination probability (**Fig. 4e**), neuronal subtype also effects myelination status, visible in the leftward shift in probability curves for neurons with highly myelinated axons (e.g. PV and VM), reaching 50% probability of myelination at just over 0.5 μm (prediction [95% confidence intervals] = VM: 0.5 μm; PV: 0.5 μm). Myelin probability curves for the remaining neuron subtypes were right shifted, reaching 50% probability of myelination at 0.6 μm (PO), 0.7 μm (RBP4), and 0.8 μm (NXPH4) (**Fig. 4e**). Together, these results indicate that myelination of cortical axons does not follow a strict diameter-myelination relationship similar to that described in the PNS^12–14^, and that other neuron intrinsic factors profoundly influence the probability of being selected for myelination.

**Figure 4.**
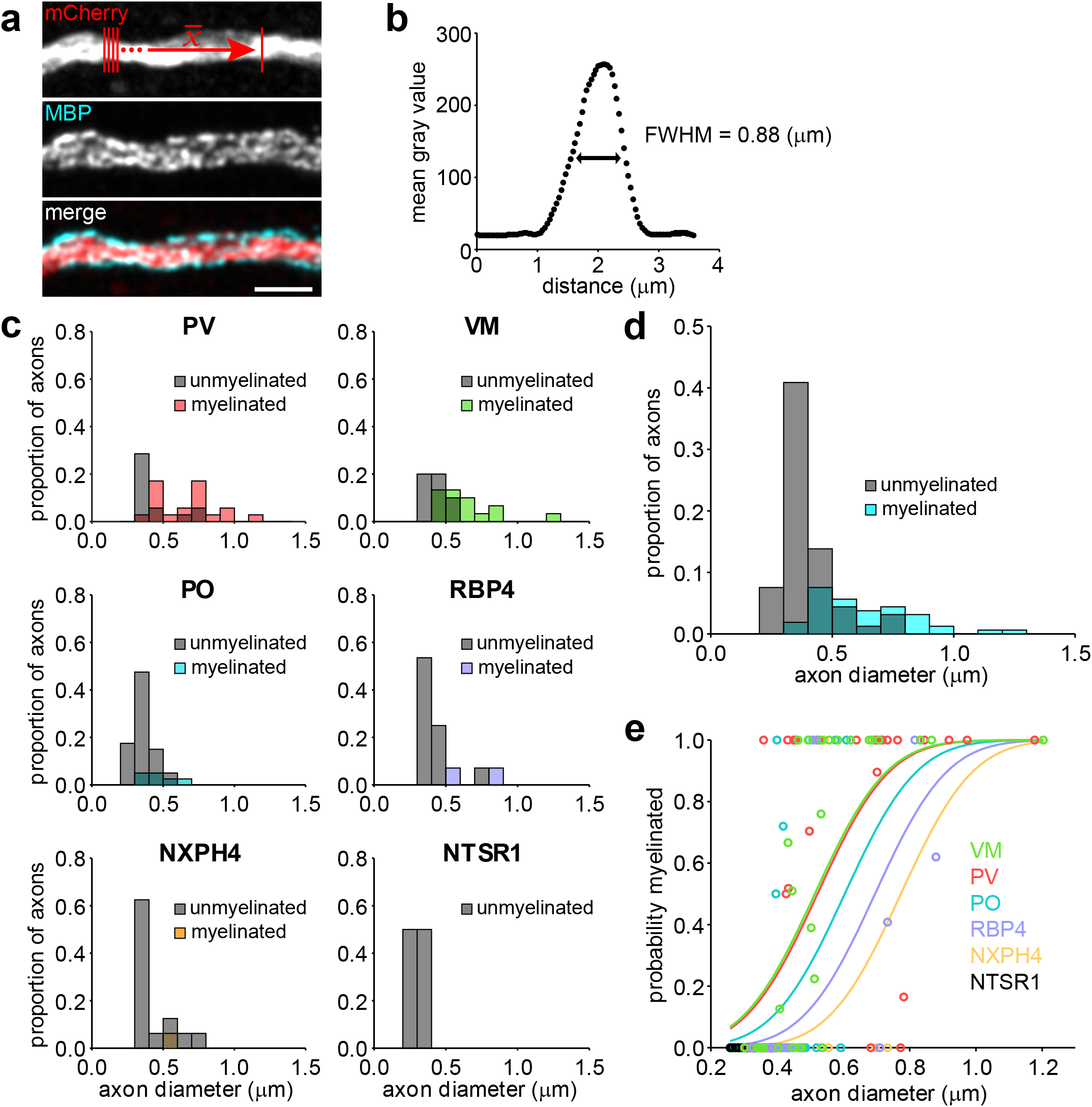
A combination of diameter and neuronal subtype predicts myelination status. **a**, Example segment of an axon imaged at high resolution for diameter analysis and corresponding MBP immunostaining. A small rectangle was used to plot a profile of the average gray value intensity profile across a few μm of axon (result shown in (**b**)) to determine its diameter by the full width at half maximum (FWHM). **c**, Plot of the average profile across the axon in (**a**). Each point represents the average gray value of a single pixel across the representative lines in (**a**), top panel. **d**, Histograms of the proportions of axons myelinated (colored bars) or unmyelinated (gray bars) for different axon diameters. Note color of overlapping bars is additive. **e**, As in (**c**), pooled across all neuronal subtypes. **f**, Probability distributions for the myelination probability of each neuronal subtype given a certain diameter. Points are individual axons myelinated. Intermediate probabilities (not 1 or 0) represent the average myelination status across at least three positions along the axon surveyed to calculate average diameter. See text for details on probability functions.

**Table 4:**
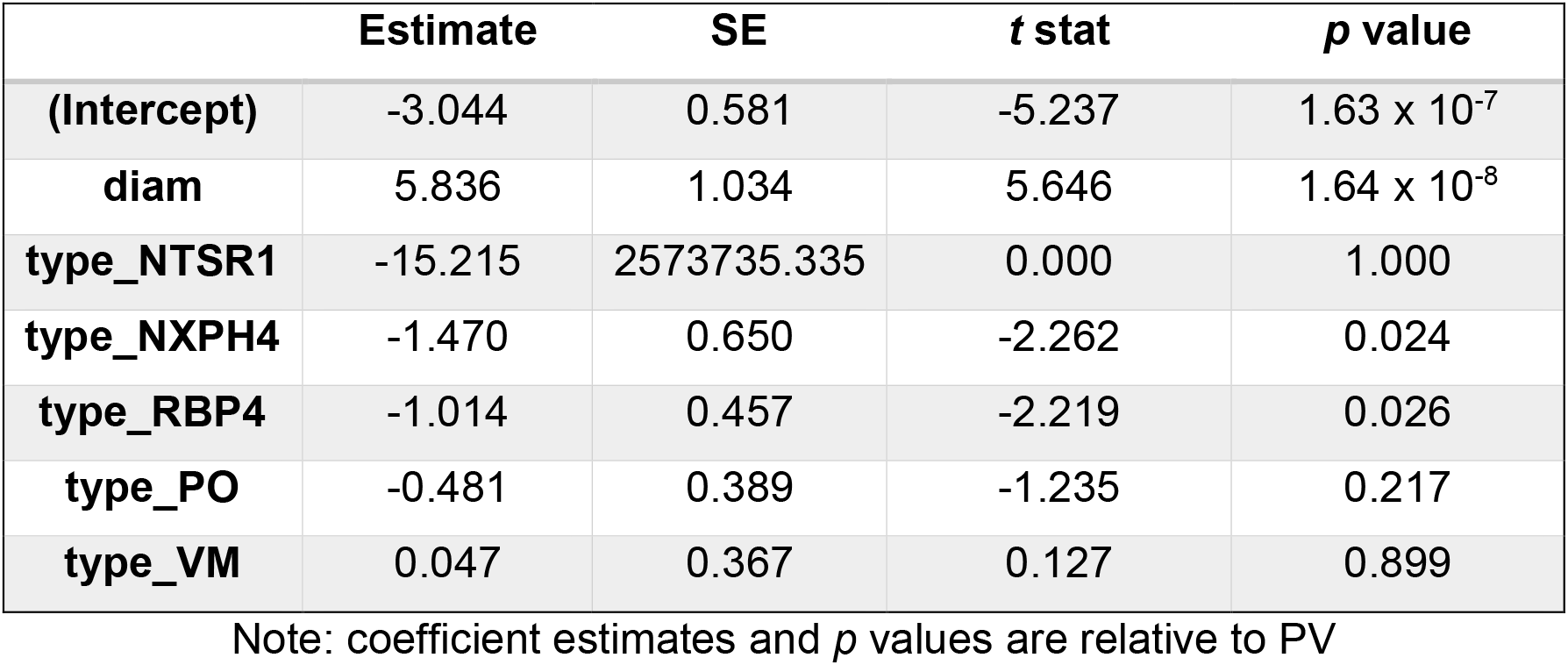
Estimated coefficients for generalized linear model

|  | **Estimate** | **SE** | **t stat** | **p value** |
| --- | --- | --- | --- | --- |
| **(Intercept)** | -3.044 | 0.581 | -5.237 | $1.63 \times 10^{-7}$ |
| **diam** | 5.836 | 1.034 | 5.646 | $1.64 \times 10^{-8}$ |
| **type_NTSR1** | -15.215 | 2573735.335 | 0.000 | 1.000 |
| **type_NXPH4** | -1.470 | 0.650 | -2.262 | 0.024 |
| **type_RBP4** | -1.014 | 0.457 | -2.219 | 0.026 |
| **type_PO** | -0.481 | 0.389 | -1.235 | 0.217 |
| **type_VM** | 0.047 | 0.367 | 0.127 | 0.899 |

Note: coefficient estimates and *p* values are relative to PV

### Remyelination restores myelin patterns among different neuronal subtypes

Recent studies indicate that oligodendrocyte regeneration from endogenous progenitors is sufficient to restore myelin levels in layer I after demyelination; however, the pattern of myelin along individual axons is altered after remyelination, with a large fraction of sheaths established on previously unmyelinated axon segments^37,53^. It is not yet known if this reorganization is equally distributed across different neuronal subtypes. If the cell intrinsic features that shape myelination during development are altered by demyelination, it could result in profound alterations in circuit properties, despite restoring myelin levels. To address this question, we used longitudinal *in vivo* imaging to compare the myelination patterns of individual PV and VM axons before and after remyelination, as they have the highest probability of being myelinated and the largest variation in myelination patterns (**Fig. 1a,b** and **Fig. 4e**). To achieve simultaneous, two color labeling of PV and VM axons *in vivo*, AAV encoding EGFP was injected into the VM nucleus of the thalamus of *PV-Cre;Ai9* mice. Cranial windows were then installed over the somatosensory barrel field before mice underwent a three-week cuprizone protocol shown previously to almost entirely demyelinate the superficial layers of cortex^37,54^. Two photon imaging was used to record the distribution of PV and VM axons in layer I and spectral confocal reflectance (SCoRe) microscopy^55^ was used to identify myelin sheaths in the same field of view. Comparison of myelin patterns before and after oligodendrocyte regeneration using this approach revealed that the majority of axons of both neuronal subtypes were myelinated before cuprizone exposure and remyelinated after five weeks of recovery (PV: 140/197 axons; VM: 150/214 axons; n = 12 regions from 9 mice) (**Fig. 5a**), indicating that the high preference for myelinating these axons is preserved after demyelination.

**Figure 5.**
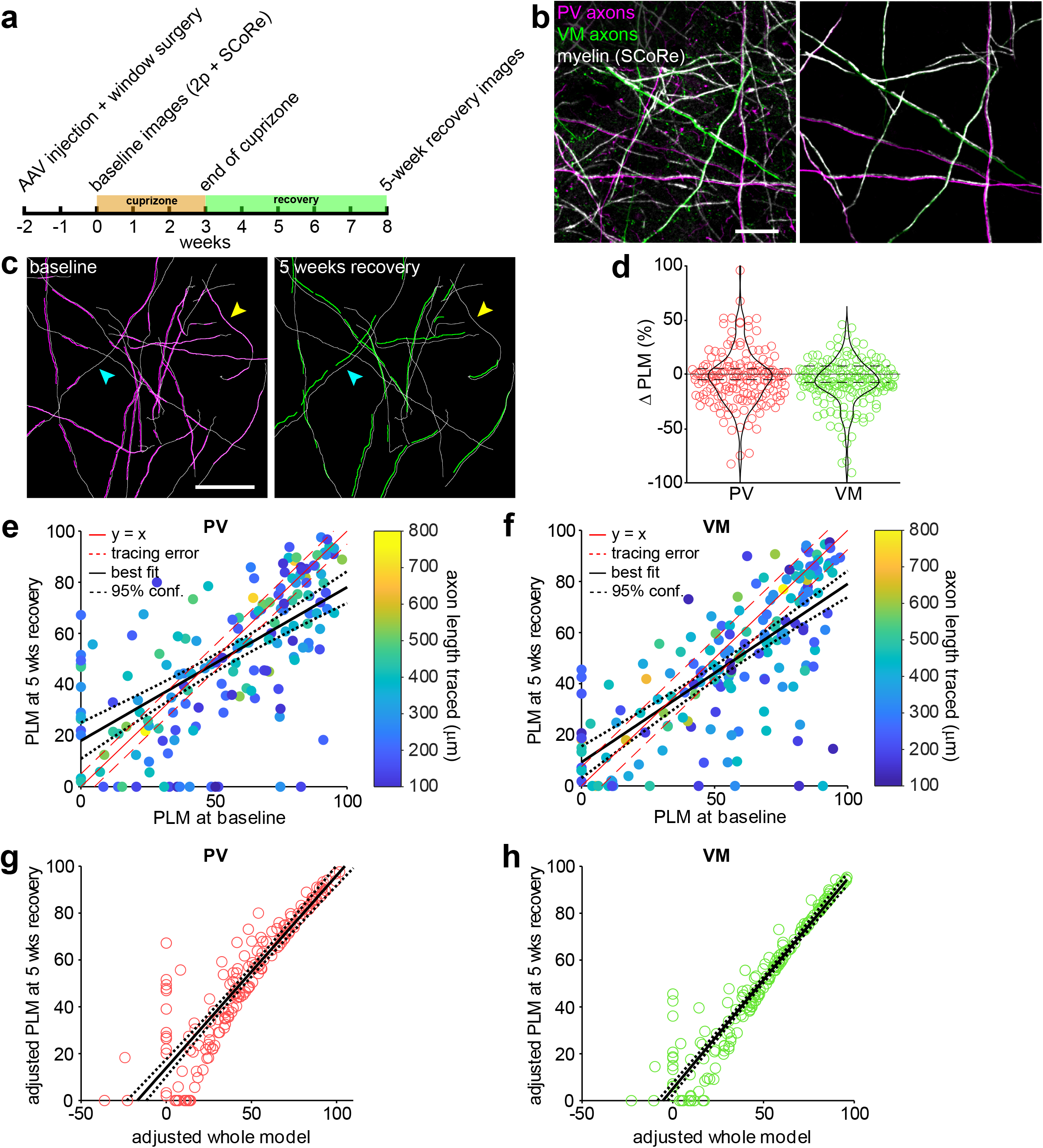
Overall subtype-specific myelination patterns are restored despite remodeling of individual axons. **a**, Experimental timeline. **b**, Example image from a *PV-Cre; Ai9* mouse injected with AAV-EGFP into thalamic VM. SCoRe signal from the same region is overlaid on top of the fluorescent dual-colored image acquired with two-photon microscopy. Right panel shows a few myelinated axons of both neuronal subtypes extracted from the left panel. **c**, Axon traces from an example three-week cuprizone timecourse. Left panel shows PV axon traces (white) and the associated myelin segments traced from the corresponding SCoRe image (magenta). Right panel shows same region with the surviving baseline axon traces overlaid with the positions of SCoRe segments traced at five weeks recovery (green). Yellow arrowhead indicates example axon that had a net loss in length myelinated, while the cyan arrowhead indicates a portion of axon which gained myelination. **d**, Pooled data from remyelinated PV and VM axons showing the net change in PLM between baseline and five weeks recovery where –100% represents an axon going from fully myelinated to fully unmyelinated, and +100% represents an axon that went from unmyelinated to fully myelinated. Dashed lines bordering 0% reflect the tracing error as calculated by the average negative Δ PLM from axons of the respective subtype traced in control animals (**Extended Data Fig. 2**, see text for additional details). **e-f**, Scatter plots showing the relationship between the PLM of an axon at baseline and its PLM at five weeks recovery from cuprizone for PV axons (**e**) and VM axons (**f**). The color of each point (an individual axon) is coded by the length of the axon traced. The solid red line is the identity line (no change between baseline and five weeks recovery) and the red dashed lines are the error based on control traces (as in (**d**)). The black line is the best fit found by regression and the black dotted lines are the 95% confidence bounds. **g-h**, Data as plotted in (**e-f**) readjusted to include the Δ PLM of each axon in the linear model (adjusted formula: PLM at 5 weeks recovery ∼ 1 + baseline PLM + baseline PLM: Δ PLM; **g**, PV axons; **h**, VM axons). Solid line is the fit and dotted lines are the 95% confidence bounds.

To determine if remyelination alters the distribution of myelin along individual PV and VM axons, each myelin internode was traced along each labeled axon within the imaging volume before cuprizone and after recovery. This analysis revealed that although the percent change in axon length myelinated (Δ PLM) for individual axons varied considerably (**Fig. 5b**), there was little change in average PLM across the population of axons for either PV or VM neurons (PV: – 3.7 ± 1.9%, VM: –7.7 ± 1.6%) (**Fig. 5c**). Together, these results indicate that while the pattern of myelination along individual axons in the cortex changes after a demyelinating event, the overall myelin content on PV and VM axons is restored by remyelination.

To define the relationship between baseline and recovery myelin patterns, we plotted the PLM for individual PV and VM axons at five weeks recovery versus baseline and fit a linear model to these distributions (**Fig. 5d,e**). This plot revealed a strong positive correlation between baseline PLM and recovery PLM for individual axons (PV: R^2^ = 0.389, *p* = 1. 01 × 10^−19^; VM: R^2^ = 0.531, *p* = 2. 86 × 10^−30^), with minimal bias from differences in the axon length traced (PV: R^2^ = 0.049, VM: R^2^ = 0.019) (**Fig. 5d,e**). Most values in these plots were close to the identity line (0% Δ PLM) at PLM values > 60%, in accordance with prior *in vivo* studies indicating that continuously-myelinated axons are most-likely to be remyelinated^37,53^. However, there were more well-myelinated axons below the identity line (solid red lines, **Fig. 5d,e**), suggesting a slight trend toward loss of myelination (Δ PLM for axons with > 75% PLM at baseline, PV: median: –6%, range: [–82%, 15%]; VM: median: –8%, range: [–90%, 10%]) (**Fig. 5c,d**).

Countering this trend, many unmyelinated axons or weakly-myelinated axons with low PLM at baseline had large gains in myelination during recovery (Δ PLM for axons with < 25% PLM at baseline, PV: median: 21%, range: [–20%, 95%]; VM: median: 6%, range: [–14%, 46%]) (**Fig. 5c,d**). As a result, the best fit lines cross the identity lines at 45% and 31% for PV and VM axons respectively, indicating that weakly myelinated axons are more likely to gain myelin during recovery, while continuously myelinated axons are more likely to lose myelin. To test this hypothesis, we refitted the linear model, testing for an interaction between PLM at baseline and the Δ PLM exhibited during recovery (recovery PLM ∼ 1 + baseline PLM + baseline PLM: Δ PLM). After adjusting for this preference, the updated model strongly predicted PLM outcomes at five weeks recovery for both PV and VM axons (PV: R^2^ = 0.773, *p* = 3.06 × 10^−54^; VM: R^2^ = 0.907, *p* = 2.87 × 10^−89^) (**Fig. 5f,g**), supporting the hypothesis that regeneration of oligodendrocytes and their myelin sheaths induces a slight shift in myelination extent from well-myelinated to sparsely-myelinated axons in both populations of neurons.

## DISCUSSION

The cerebral cortex contains axons that arise from the local collaterals and long-range projections of a diverse population of neurons. Despite the presence of axons of suitable size and the persistence of oligodendrocyte progenitors, myelin density in the cortex remains low. The diverse patterns of myelin observed along cortical axons, even among axons from distinct neuron subtypes, raise many new questions about the mechanisms that establish these patterns and the impact of this complex organization on myelin repair. Our *in vivo* results reaffirm that axonal diameter is a key factor in determining myelination status in this region, as neurons that had the largest axon diameters also had the highest myelin content, recapitulating *in vitro* studies showing that cortical oligodendrocytes rarely form wraps around structures less than 0.4 µm^15,56^, and the wealth of anatomical studies in the CNS showing that thinner axons are less likely to be myelinated. Nevertheless, many PV and VM axons of relatively thin diameter (< 0.5 µm) were continuously myelinated in layer I of the somatosensory cortex, and many large diameter axons (> 0.5 µm) were unmyelinated. Moreover, unmyelinated segments often occurred immediately adjacent to myelinated regions along axons with consistent diameter (**Fig. 2f**). The lower rate of oligodendrogenesis in the cortex relative to white matter^57,58^, and the strict, cell intrinsic limits on myelin output by individual oligodendrocytes^15,19,37^, may allow such discontinuous patterns to persist despite the high preference of these axons for myelination. Although oligodendrogenesis and myelination continues in the mouse cortex for many months after birth^9,10,57–59^, the rate of oligodendrogenesis in these five month-old mice is very low (< 1 new oligodendrocyte per month within the imaging volume)^37,58,60,61^, and individual myelin sheaths are extremely stable^37^, indicating that the diverse myelination patterns observed on these distinct excitatory and inhibitory neurons are compatible with the advanced processing capabilities of cortical circuits in the adult brain.

### Regulation of cortical axon myelination

Although axon diameter is a key determinant of myelination probability, recent evidence suggests that target selection by oligodendrocytes is also influenced by the frequency of axon collateral formation^52^, with extensively branched terminal arbors less likely to be myelinated; thus, differences in axonal branching may also contribute to the range of myelination profiles observed in layer I. Consistent with this hypothesis, thalamocortical PO axons that exhibit more terminal arborizations in the somatosensory cortex were less likely to be myelinated for a given axon diameter than VM axons (**Fig. 4c**), which have more long-range projections that “pass through” the somatosensory region with limited branching (**Extended Data Fig. 1i-l**).

In addition to these morphological constraints, our results reveal that neurons exert cell autonomous control of myelination independent of the diameter and degree of branching of their axons. This finding is consistent with the observation that oligodendrocytes do not form myelin on dendrites, cell bodies, or capillaries, structures that by size alone should be permissive targets. At present, we know very little about the molecular mechanisms that regulate target selection by oligodendrocytes. Recent results indicate that integral membrane proteins (e.g. JAM2, GPCRs) that participate in cell adhesion can both promote or prevent myelination^25,62^, suggesting that adhesive and repulsive interactions could bias this selection process by regulating extension of lamellar sheaths from nascent oligodendrocyte processes^63,64^, with the relative amounts of these proteins accounting for the bias in subtype remyelination observed. As neuronal activity can promote both oligodendrogenesis and myelination of some neurons^20–22,65– 67^, it is possible that activity adjusts myelination probability by altering the expression, trafficking or localization of these proteins

The vast territory available for addition of new myelin on axons that have a high probability of being myelinated (e.g. large diameter, low branching) and the persistence of oligodendrocyte progenitors creates opportunities for circuit modification in adulthood by increasing myelin content, a phenomenon termed adaptive myelination. Indeed, motor learning, exposure to new environments, and other life experiences correlate with changes in oligodendrogenesis, myelination and altered network activity^68^. However, the mechanisms underlying myelin plasticity are unknown and the impact of these changes on circuit behavior remain largely theoretical^69^. *In vivo* imaging studies have revealed that less than half of newly formed oligodendrocytes in the cortex become stably integrated^70^, suggesting that shifts in myelin content could be achieved by simply regulating integration probability rather than promoting oligodendrogenesis. It is not yet known what types of neurons increase their myelin content in these contexts, whether such changes are long lasting or how such changes ultimately influence the properties of cortical circuits. It is also unclear what adaptive advantage is provided by the gradual addition of myelin during adulthood. The ability to monitor the myelination patterns along individual axons of genetically distinct neuronal subtypes using longitudinal *in vivo* imaging may provide the means to answer these questions.

### Restoration of cortical myelin through oligodendrocyte regeneration

The goal of regenerative processes is to restore function. The extraordinary stability of individual myelin sheaths, in terms of their position and length along cortical axons suggests that these patterns are important for axonal output and thus overall circuit function. However, prior studies demonstrated that overall myelin patterns change in the cortex after destruction and regeneration of oligodendrocytes^37,53,54^, and we show here that the position of internodes and total myelin content along individual axons of both local inhibitory PV interneurons and long-range excitatory VM thalamocortical neurons change substantially after remyelination. Nevertheless, following the loss and regeneration of myelin sheaths along individual axons of defined neuron subtypes *in vivo* revealed that the net change myelin content (Δ PLM) across both populations of axons was negligible after regeneration, despite the considerable reorganization of myelin along individual axon segments. These findings indicate that the preference for myelinating these axons is preserved after demyelination and raise the possibility that oligodendrocyte regeneration is optimized to restore overall myelin content across the population of diverse axons, rather than the precise patterns of myelin along individual axons. This hypothesis is consistent with the opportunistic behavior of oligodendrocytes, in which axonal selection is determined by a hierarchy of signals (axon diameter, axonal branching, cell identity, activity) that define the myelination probability of an individual axon. As oligodendrocytes are regenerated in new locations^37^, reorganization of exact myelin patterns is precluded, but overall myelin content can be preserved. It is important to note that many of the axons traced in this study extended outside of the bounds of the imaged volumes. Thus, it is possible that the axons with large changes in PLM experienced equal and opposite changes outside the area examined, resulting in precise restoration of myelin content (though not position) along individual axons. New advances that allow imaging over larger areas of the cortex^71^ and at greater depths (three photon excitation) may enable myelin to be visualized on individual axons across several millimeters of cortex to resolve this question.

PV and VM axons are the most highly myelinated axons investigated in this study; however, due to the high density and vast diversity of axons in layer I, the myelin on these two axon populations represents less than half of the myelin in this region. While we did not determine the remyelination efficiency of other less myelinated neuronal subtypes, similar outcomes are likely. Previous remyelination studies in layer I revealed that isolated sheaths on discontinuously myelinated axons were rarely replaced^37,53^ and the number of isolated sheaths lost was approximately the same as those gained on previously unmyelinated axon segments^37^. If myelination patterns were strongly altered in these weakly-myelinated axon populations, we would have expected the net Δ PLM to be significantly below zero, as myelin is “transferred” to other weakly myelinated populations by regenerated oligodendrocytes.

The functional implications of these changes in myelination patterns in the cerebral cortex are unknown. Cuprizone-induced demyelination has been shown to cause pyramidal neuron hyperexcitability by shifting axon initial segment position, causing slowed, continuous, rather than saltatory conduction of action potentials and increased conduction failure^72,73^. Moreover, inflammatory myelin loss (EAE, experimental autoimmune encephalomyelitis) results in a dramatic loss of excitatory synapses in both mouse and primate^74^. Hence, alterations in cortical myelin patterns could not only alter action potential conduction and network oscillations, but also neuronal connectivity and synaptic plasticity. However, neural circuits are capable of extraordinary adaptation and it is possible that restoring the overall myelin content across a population of axons, without precise replacement of individual internodes, is sufficient to restore circuit function. This possibility provides hope for therapeutic strategies for treating MS based on global promotion of oligodendrogenesis^54,75–77^, as the intrinsic properties of axons and the opportunistic behavior of oligodendrocytes may help reestablish the distinct myelin profiles of different neurons in the cortex.

## METHODS

### Animal care and use

Female and male adult mice were used for experiments and randomly assigned to experimental groups. All mice were healthy and did not display any overt behavioral phenotypes. Mice were maintained on a 12-h light/dark cycle, housed in groups no larger than 5, and food and water were provided *ad libitum* (except during cuprizone-administration, see below). All animal experiments were performed in strict accordance with protocols approved by the Animal Care and Use Committee at Johns Hopkins University. The following transgenic mouse lines were used in this study:

Pvalb-IRES-Cre*^78^

Sst-IRES-Cre^78^

GN220-Ntsr1-Cre^45^

Rbp4-KL100^45^

Nxph4-2A-CreERT2-D^46^

Rosa-CAG-LSL-tdTomato-WPRE (Ai9)*^44^

Rosa-CAG-LSL-EYFP-WPRE (Ai3)^44^

*note: C57BL/6 congenic strains were used for cuprizone experiments

For axon tracing in immunostained flatmounts, mice were aged to five months, when cortical myelination has nearly plateaued^59,61^.

### CreER induction

To induce tdTomato expression in the NXPH4-CreER mice, five-month old mice were injected with 4-hydroxytamoxifen (4-HT, Sigma) dissolved in sunflower seed oil (Sigma) and administered by i.p. injection twice daily for five days at a dose of 100 mg/kg bodyweight. Mice were perfused two weeks from the last day of tamoxifen injection.

### Viral injections

Five-month old mice were anesthetized with isoflurane (induction, 5%, mixed with 1 L/min O_2_; maintenance, 1.5–2%, mixed with 0.5 L/min O_2_), and their scalps shaved with an electric trimmer. Mice were transferred to a homeostatic heating pad set to 37°C and their heads secured by ear bars on an Angle 2 stereotax. An incision was performed along the midline of the scalp to expose the skull. Bregma was identified and zeroed in the Angle 2 software. The skull was leveled by ensuring z measurements 2 mm left and right from bregma differed by < 0.01 mm and difference in z between bregma and lambda was < 0.1 mm. A small craniotomy was drilled directly above the target coordinates for the injection (VM: −1.45 mm AP, 0.80 mm ML, −4.25 mm DV; PO: −1.35 mm AP, 1.23 mm ML, −3.30 mm DV; MOp: 1.35 mm AP, 1.5 mm ML, −1.83 mm DV) and a micropipette filled with virus (AAV-DJ-CaMKii-mCherry, 1.2 × 10^13^, or AAV9-hSyn-EGFP-WPRE-bGH, at a titer of 4.44 × 10^11^, or AAV9-CAG-FLEX-tdTomato-WPRE-bGH, at a titer of 1.55 × 10^11^) was slowly lowered to the injection site. Virus was delivered in 13.8 nL pulses over the course of 15 minutes for a total volume of 150 to 250 nL. After a five-to ten-minute wait period to allow diffusion, the pipette was slowly withdrawn, the craniotomy sealed with VetBond, and the scalp closed with metal clamps. For subsequent histological preparation, mice recovered for two weeks to allow sufficient expression and then perfused. For subsequent *in vivo* imaging, cranial windows were installed immediately following the injection (best window clearing rates) or four to five days later (reduced window clearing rates).

### Flatmount preparation

Mice were deeply anesthetized with sodium pentobarbital (100 mg/kg w/w) and perfused transcardially first with 20-25 mL warm (∼30° C) PBS (pH 7.4) followed by 20-25 mL ice-cold 4% paraformaldehyde (PFA in 0.1 M phosphate buffer, pH 7.4). Brain hemispheres not used for flatmounts were then post-fixed in 4% PFA for 12 hours, before being transferred to a 30% sucrose solution (in PBS, pH 7.4). To generate cortical flatmounts, cortical mantles were carefully dissected from underlying structures directly after perfusion and pressed between glass slides separated by ∼1 mm. Cortices were postfixed in this position in 4% PFA for 6 to 12 hours at 4° C before being unclamped from the glass slides and transferred to 30% sucrose solution. Tissue was stored at 4° C for more than 48 h before sectioning. Brain tissue was frozen in TissueTek and sectioned at 35 to 50 μm thickness on a cryostat (Thermo Scientific Microm HM 550) at −20° C. For flatmount sections, extreme care was taken to ensure the cutting plane was perfectly horizontal to the flatmount surface in order to ensure layer I was contained within the section. First, a mounting chuck was covered with TissueTek and frozen. The mound of TissueTek was then cut until a flat surface across the diameter of the chuck was achieved. Flatmounts were placed pia-down onto a silanized glass slide (hydrophobicity helps prevent tissue sticking), covered with TissueTek, and flipped onto the pre-cut mound. After the tissue was completely frozen, the glass slide was removed, the edges of the TissueTek mound were beveled, and the chuck was mounted for sectioning.

### Immunohistochemistry

Immunohistochemistry was performed on free-floating sections. Sections were preincubated in blocking solution (5% normal donkey serum, 0.3% Triton X-100 in PBS, pH 7.4) for 1 or 2 hours at room temperature, then incubated for 24 to 48 hours at 4°C or room temperature in primary antibody (listed in Key Resources Table). Secondary antibody (see Key Resources Table) incubation was performed at room temperature for 2 to 4 hours or overnight at 4° C. Sections were mounted on slides with Aqua Polymount (Polysciences).

### Cranial windows

Cranial windows were prepared as previously described^37^. Briefly, mice 8 to 10 weeks old were anesthetized with isoflurane (induction, 5%, mixed with 1 L/min O_2_; maintenance, 1.5–2%, mixed with 0.5 L/min O_2_), and their body temperature was maintained at 37° C with a thermostat-controlled heating plate. The scalp over the right hemisphere was removed and the skull cleaned and dried. A custom metal head plate with a central hole was attached to the skull with dental cement (C and B Metabond) for head stabilization. The head plate was then fixed in place by clamping head bars while a three-millimeter diameter circular region of skull over somatosensory cortex (–1.5 mm posterior and 3.5 mm lateral from bregma) was removed using a high-speed dental drill. A piece of cover glass (VWR, No. 1) was placed in the craniotomy and sealed with cyanoacrylate glue (VetBond (3M) and Krazy Glue).

### Simultaneous *in vivo* imaging of PV and VM remyelination

Baseline images were acquired two to three weeks after cranial window installation, and mice with clear windows were randomly assigned to cuprizone or control conditions. Due to the rarity of animals having the correct genotype, successful intracranial viral injections, and clear windows, some animals with bone regrowth under the window at two to three weeks after the cranial window surgery had their windows repaired. Cyanoacrylate glue holding the original coverslip was carefully drilled away, the coverslip was removed, and invading bone into the window region was cut away before gluing in a new glass coverslip. Mice with repaired windows recovered for an additional week before checking the window clarity and taking baseline images. This procedure was performed for mice assigned to both control and cuprizone-treated groups.

During each imaging session (at baseline, one-week recovery, and five-weeks recovery), mice were anesthetized with isoflurane (induction, 5%; maintenance, 1.5–2%, mixed with 0.5 L/min O_2_ and fixed by their headplates in a custom stage. Two-photon images were collected using a Zeiss LSM 710 microscope equipped with a GaAsP detector using a mode-locked Ti:sapphire laser (Coherent Ultra) tuned to 1000 nm. The average power at the sample during imaging was < 30 mW. Vascular landmarks and axon orientations were used to identify the same cortical area across imaging sessions. Z stacks were 425 μm x 425 μm x 110 μm acquired at a resolution of 2048 × 2048 pixels using a coverslip-corrected Zeiss 20x water-immersion objective (NA 1.0).

Immediately after collecting the two-photon image, the sample was imaged again in reflectance mode to collect SCoRe signal with a z stack of the same dimensions as the two-photon image. SCoRe has been shown by several groups independently to faithfully detect myelin sheaths in layer I of the cortex^9,10,54^. Laser parameters for reflectance imaging were used as previously described^9^, and the pinhole was set to 1.5 Airy units.

Comparing SCoRe signal between baseline and recovery without visualization of oligodendrocyte cell bodies and processes meant surviving sheaths and sheaths lost and replaced in the same position were unable to be differentiated. While more frequent imaging may have allowed us to detect timepoints in which SCoRe signal was absent before reappearing in a subsequent timepoint, we found that weekly imaging of two-photon and SCoRe signal produced phototoxicity and increased the rate of axon damage. Our previous experiments using a three-week cuprizone model found an average of 15.6% of all sheaths survive cuprizone in layer I^37^. Assuming these surviving sheaths would be randomly distributed across axon types, we estimate that surviving sheaths represent a very small fraction of the myelin analyzed and that the majority of overlapping SCoRe signal between baseline and five-weeks recovery is remyelinated.

### Cuprizone administration

At 9 to 11 weeks of age, male and female *Mobp-EGFP* or C57BL/6 mice were fed a diet of milled, irradiated 18% protein rodent diet (Teklad Global) alone (control) or supplemented with 0.2% w/w bis(cyclohexanone) oxaldihydrazone (Cuprizone, Sigma-Aldrich) in gravity-fed food dispensers for three weeks. Both control and cuprizone-treated mice were returned to a regular pellet diet after three weeks during the recovery period^37^.

### Image collection

Images were acquired using a confocal laser-scanning microscope (Zeiss LSM 510 Meta; Zeiss LSM 710; Zeiss LSM 880). For population analyses (Figure 1), 5 × 5 tiled z stacks (650 μm x 650 μm x 40 μm, z slice thickness: 0.5 μm, pinhole set to 1 Airy unit for each wavelength, 2048 × 2048 pixels) were acquired with a Zeiss 63x oil objective (NA 1.4) in primary somatosensory cortex in a cortical flatmount. Regional comparisons for PV axons used 2 × 2 tiled z stacks acquired at 63x. For diameter analysis, individual z stacks were acquired at 63x in Zeiss Airyscan mode (59.65 μm x 59.65 μm or 78.01 μm x 78.01 μm, 2048 × 2048 pixels, z slice thickness: 0.21 μm, pinhole: 1 Airy unit). Tiled overviews of flatmounts and coronal sections were acquired at 4x with a Keyence epifluorescence microscope.

### Diameter calculations

Within high resolution z stacks of each neuronal subtype, the distribution of gray values across the width of individual axons were averaged across a 2-μm length (ImageJ “Plot Profile”), and the full-width at half maximum (FWHM) was used to represent the diameter. Two to three of these measurements were taken for a 35-μm segment of axon within the imaging volume and averaged together for each individual axon. The probability myelinated for each axon was calculated as the number of measurement locations that were unmyelinated divided by the number of measurement locations that were myelinated. Given that the very small image area required for sufficient resolution (see **Image collection**) was close to the average length of myelin internodes, the majority of axons had probability values of either 0 or 1.

### Axon and myelin tracing

In all experiments, channels containing axon and myelin information were first split before tracing began to blind experimenters to myelination status of each axon. Axons were selected for tracing by placing a 100 μm x 100 μm grid across the image and a random number generator selected grid coordinates for each axon seed. The first axon from the pia observed reaching a length of at least 100 μm (passed across the grid square) was selected and traced from that grid square along its full length within the image in both directions. Axon branches within the image were also traced if their lengths were also at least 100 μm in length. Axon traces were then imported into the myelin channel (either MBP immunolabeling or SCoRe signal) and used to trace myelin segments associated with each axon.

Fluorescently-labeled axons were traced in Fiji using Simple Neurite Tracer/SNT^79^. Traced segments were put through a smoothing function prior to length calculations to reduce artifacts of jagged traces. As axons and their associated MBP segments were traced in their respective channels and signal properties differed between them, some continuously-myelinated axons (completely covered by myelin save for nodes of Ranvier) had PLM values > 100% if jitter in MBP traces was higher than the axon traces leading to an artefactually higher total sheath length than total axon length. In these cases, PLM was manually set to 99%.

### Image processing and analysis

Image stacks and time series were analyzed using Fiji. Images used in figures were adjusted for brightness and contrast levels for clarity. *In vivo* z stacks were de-noised with a 3D median filter (1-pixel radius). SCoRe images were first background subtracted (15-pixel rolling ball), 3D medial filtered (1-pixel radius), and all three channels were summed together for a final SCoRe channel used for tracing. Because virally expressed EGFP signal from VM thalamocortical axons strongly bled through into the red channel at 1000 nm excitation, the green channel was subtracted from the red channel before tracing tdTomato-positive PV axons. Longitudinal image stacks were randomized for analysis and traced by a blinded experimenter and revised by a second blinded experimenter. Baseline and five-week recovery timepoints were randomly assigned as “A” or “B” and experimenters always made original axon traces using the “A” timepoint. In some cases when baseline was the “A” timepoint, traced axons were no longer visible in the recovery timepoint (either by part of the imaged region being obscured by meningeal thickening/bone growth or by axonal degeneration, which occurred at low rates in both control (VM: 8 ± 5%, PV: 1 ± 1%) and cuprizone-treated (VM:14 ± 3%, PV: 3 ± 1%) animals). These rates of loss were not significantly different between control and cuprizone conditions (VM: p = 0.38, PV: p = 0.46, two-sample unpaired t-test). In these cases, missing axons were excluded from analysis. Axons which remained unmyelinated at both timepoints (PLM = 0) were not included in linear models so as not to strongly bias trends of myelin replacement.

### Statistical analysis

Statistical analyses were performed with MATLAB (Mathworks). Significance was typically determined using Kruskal-Wallis one-way ANOVA with Dunn–Šidák correction for multiple comparisons. Violin plots were created using violin.m^80^. Each figure legend or the text otherwise contains the number of animals used, statistical tests used to measure significance, and the corresponding significance level (*p* value). Data are reported as mean ± standard error, or prediction [lower 95% confidence bound, upper 95% confidence bound], and *p* < 0.05 was considered statistically significant.

## Data availability

All published code, tools, and reagents will be shared on an unrestricted basis; requests should be directed to the corresponding authors. MATLAB scripts and ImageJ macros are available at https://github.com/clcall/Call_Bergles_2021_CTSM.

## ACKNOWLEDGEMENTS

We thank Dr. S P Brown and previous lab members for generously providing Cre lines used in this study, Dr. M Pucak for technical assistance and impeccable instrument maintenance, T Shelly for machining expertise, T Bhardwaj and J H Kim for assistance in axon tracing, J Orthmann-Murphy for assistance in cranial window surgeries, and members of the Bergles laboratory for insightful discussions. C Call was supported by a National Science Foundation Graduate Research Fellowship (DGE-1746891). Funding was provided by grants from the NIH (NS051509, NS050274, NS080153), and the Dr. Miriam and Sheldon G Adelson Medical Research Foundation to D Bergles.

## AUTHOR CONTRIBUTIONS

CLC conceived of and designed experiments, collected data, performed analyses and wrote the manuscript. DEB conceived of and designed experiments and wrote the manuscript.

## EXTENDED DATA FIGURE LEGENDS

**Extended Data Figure 1.**
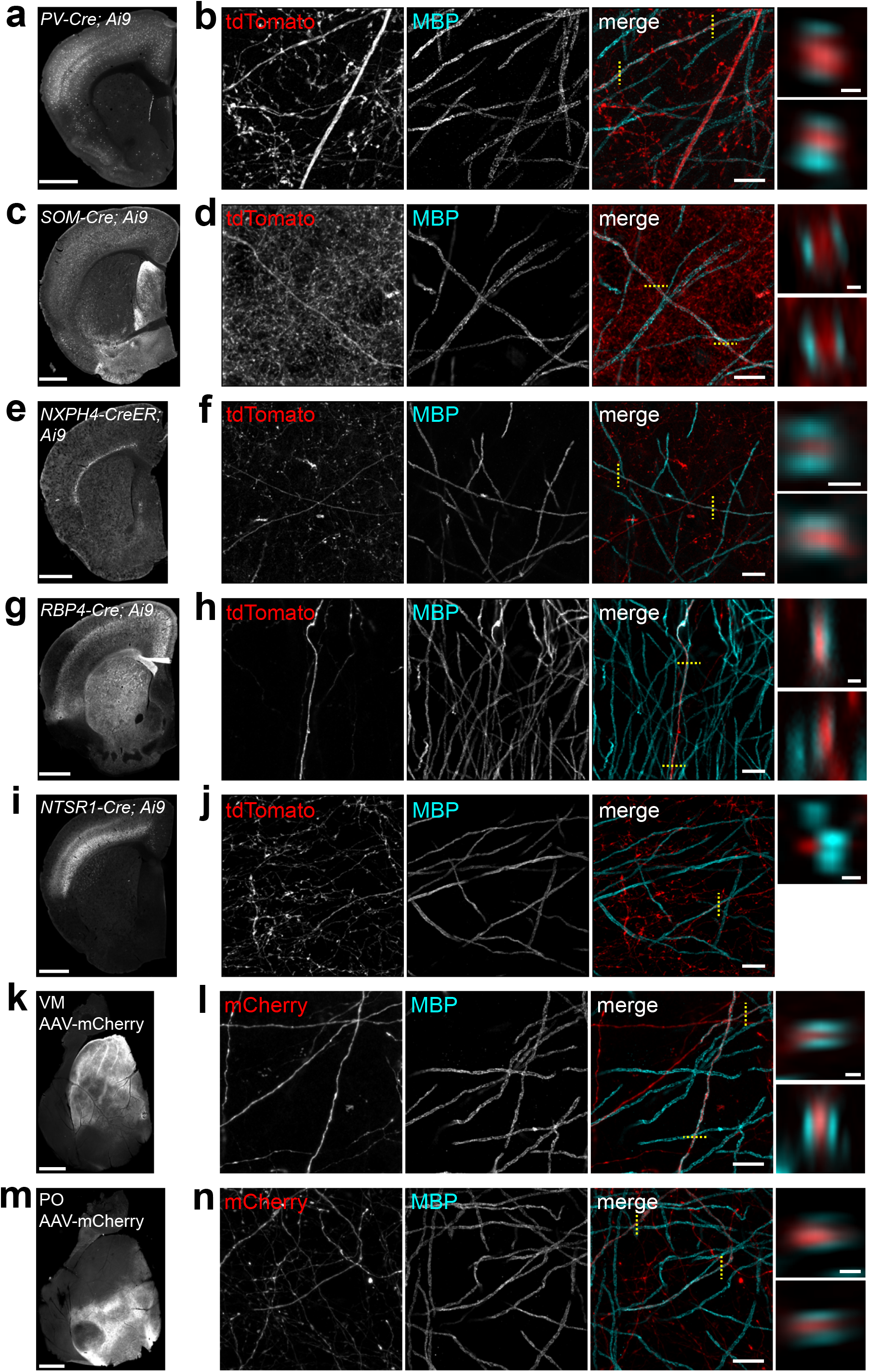
Evidence for myelination in neuron specific Cre lines and virally labeled axon populations. **a–j**, Coronal hemisections immunolabeled with MBP from brains expressing tdTomato by various Cre lines used in this study and corresponding high-resolution confocal images (< 10 μm maximum z projections) highlighting evidence for myelinated axons. **a–b**, PV-Cre. **c–d**, SOM-Cre. **e–f**, NXPH4-CreER. **g–h**, RBP4-Cre. **i–j**, NTSR1-Cre. Flatmounts and corresponding images of myelinated axons expressing mCherry from AAV injection into the ventromedial (VM, **k–l**), or posterior (PO, **m–n**), nucleus of thalamus. xz or yz slices on the far right correspond to each dotted yellow line in the merged channel images showing optical cross sections of MBP signal surrounding tdTomato- or mCherry-expressing axons (or in **j**, the lack of ensheathment). Scale bars from left to right, 1 mm, 10 μm, 0.5 μm.

**Extended Data Figure 2.**
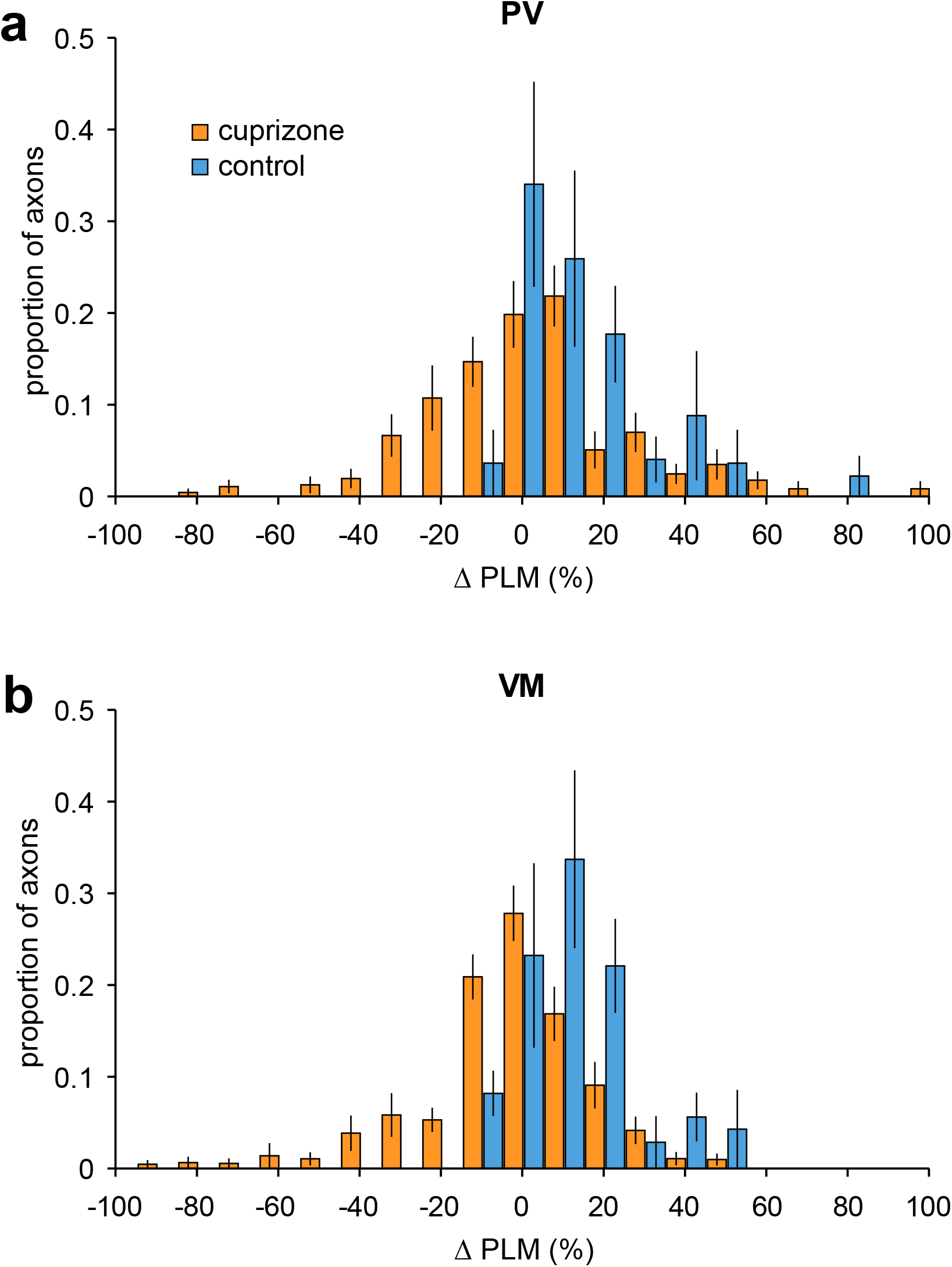
Distributions of cuprizone and control Δ PLM. **a-b**, Histograms of the average changes in PLM for PV (**a**) and VM (**b**) axons traced in mice treated with cuprizone (orange) and in control mice (blue). The control axon distributions are significantly skewed right (few axons showed negative Δ PLM), p = 0.03 (for both PV and VM), two-sample Kolmogorov-Smirnov test. There is no difference in the distributions between cuprizone treated PV and VM axons (p = 0.77 two-sample Kolmogorov-Smirnov test).

**Supplementary Video 1** – Methodology of axon and myelin tracing.

Flatmounts containing fluorescent subtype-specific axons were immunostained for MBP. Confocal tiled z stacks (675 µm x 675 µm x 40 µm) were acquired at 63x and stitched together. Channels were split and individual axons within the volume were traced (green line) blinded to their myelination status. These axon traces were subsequently loaded into the MBP channel to find myelin sheaths associated with each traced axon.

## TABLES

### Key Resources

#### Primary antibodies

| **Target Protein/markers** | **Host species** | **Source** | **Catalog #** | **Identifier** | **Additional information** |
| --- | --- | --- | --- | --- | --- |
| GFP | Chicken | Aves Lab | GFP-1020 | RRID:AB_2307313 | 1:4000 |
| mCherry | Goat | SicGen | AB0040 | RRID:AB_2333092 | 1:5000 |
| MBP | Mouse | Sternberger | 808401 | RRID:AB_2564741 | 1:2000 |
| MBP | Chicken | Aves Lab | F-1005 | RRID:AB_2313550 | 1:500 |

#### Secondary antibodies

| Target Species | Conjugate | Source | Dilution | Catalog # | Identifier |
| --- | --- | --- | --- | --- | --- |
| Anti-Mouse | Cy5 | Jackson Immuno | 1:2000 | 715-175-151 | RRID:AB_2340820 |
| Anti-Goat | Cy3 | Jackson Immuno | 1:2000 | 705-166-147 | RRID:AB_2340413 |
| Anti-Chicken | Alexa 488 | Jackson Immuno | 1:2000 | 703-546-155 | RRID:AB_2340376 |
| Anti-Chicken | Cy5 | Jackson Immuno | 1:2000 | 703-006-155 | RRID:AB_2340347 |

#### Software and Algorithms

| Name | Source | Identifier |
| --- | --- | --- |
| ZEN Blue/Black | Zeiss | RRID:SCR_013672 |
| Fiji | http://fiji.sc | RRID:SCR_002285 |
| Simple Neurite Tracer and SNT | https://imagej.net/SNT | RRID:SCR_016566 |
| Adobe Illustrator CS4 | Adobe | RRID:SCR_014198 |
| MATLAB | Mathworks | RRID:SCR_001622 |

## References

1. Cauller, L. Layer I of primary sensory neocortex: where top-down converges upon bottom-up. Behav. Brain Res. 71, 163–170 (1995).

2. Xu, N.-L. et al. Nonlinear dendritic integration of sensory and motor input during an active sensing task. (2012) doi:10.1038/nature11601.

3. Cruikshank, S. J. et al. Thalamic Control of Layer 1 Circuits in Prefrontal Cortex. J. Neurosci. 32, 17813–17823 (2012).

4. Cauller, L. J., Clancy, B. & Connors, B. W. Backward cortical projections to primary somatosensory cortex in rats extend long horizontal axons in layer I. J. Comp. Neurol. 390, 297–310 (1998).

5. Rubio-Garrido, P., Pérez-De-Manzo, F., Porrero, C., Galazo, M. J. & Clascá, F. Thalamic input to distal apical dendrites in neocortical layer 1 is massive and highly convergent. Cereb. Cortex 19, 2380–2395 (2009).

6. Zilles, K., Palomero-Gallagher, N. & Amunts, K. Myeloarchitecture and Maps of the Cerebral Cortex. in Brain Mapping: An Encyclopedic Reference vol. 2 137–156 (Elsevier Inc., 2015).

7. Micheva, K. D. et al. A large fraction of neocortical myelin ensheathes axons of local inhibitory neurons. Elife 5, 1–29 (2016).

8. Stedehouder, J. et al. Fast-spiking Parvalbumin Interneurons are Frequently Myelinated in the Cerebral Cortex of Mice and Humans. Cereb. Cortex 27, 5001–5013 (2017).

9. Hill, R. A., Li, A. M. & Grutzendler, J. Lifelong cortical myelin plasticity and age-related degeneration in the live mammalian brain. Nat. Neurosci. 21, 683–695 (2018).

10. Hughes, E. G., Orthmann-Murphy, J. L., Langseth, A. J. & Bergles, D. E. Myelin remodeling through experience-dependent oligodendrogenesis in the adult somatosensory cortex. Nat. Neurosci. 21, 696–706 (2018).

11. Tomassy, G. S. et al. Distinct profiles of myelin distribution along single axons of pyramidal neurons in the neocortex. Science (80-.). 344, 319–324 (2014).

12. Voyvodic, J. T. Target size regulates calibre and myelination of sympathetic axons. Nature (1989) doi:10.1038/342430a0.

13. Elder, G. a., Friedrich, V. L. & Lazzarini, R. a. Schwann cells and oligodendrocytes read distinct signals in establishing myelin sheath thickness. J. Neurosci. Res. 65, 493–499 (2001).

14. Schröder, J. M., Bohl, J. & Brodda, K. Changes of the ratio between myelin thickness and axon diameter in the human developing sural nerve. Acta Neuropathol. 43, 169–178 (1978).

15. Bechler, M. E., Byrne, L. & Ffrench-Constant, C. CNS Myelin Sheath Lengths Are an Intrinsic Property of Oligodendrocytes. Curr. Biol. 25, 2411–2416 (2015).

16. Mayoral, S. R., Etxeberria, A., Shen, Y.-A. A. & Chan, J. R. Initiation of CNS Myelination in the Optic Nerve Is Dependent on Axon Caliber. Cell Rep. 25, 544-550.e3 (2018).

17. Remahl, S. & Hildebrand, C. Changing relation between onset of myelination and axon diameter range in developing feline white matter. J. Neurol. Sci. 54, 33–45 (1982).

18. Waxman, S. G. & Sims, T. J. Specificity in central myelination: evidence for local regulation of myelin thickness. Brain Res. 292, 179–185 (1984).

19. Chong, S. Y. C. et al. Neurite outgrowth inhibitor Nogo-A establishes spatial segregation and extent of oligodendrocyte myelination. Proc. Natl. Acad. Sci. 109, 1299–1304 (2012).

20. Demerens, C. et al. Induction of myelination in the central nervous system by electrical activity. Proc. Natl. Acad. Sci. 93, 9887–9892 (1996).

21. Stevens, B., Porta, S., Haak, L. L., Gallo, V. & Fields, R. D. Adenosine: A Neuron-Glial Transmitter Promoting Myelination in the CNS in Response to Action Potentials. Neuron 36, 855–868 (2002).

22. Gibson, E. M. et al. Neuronal activity promotes oligodendrogenesis and adaptive myelination in the mammalian brain. Science 344, (2014).

23. Wake, H., Lee, P. R. & Fields, R. D. Control of local protein synthesis and initial events in myelination by action potentials. Science 333, 1647–1651 (2011).

24. Wake, H. et al. Nonsynaptic junctions on myelinating glia promote preferential myelination of electrically active axons. Nat. Commun. 6, 7844 (2015).

25. Redmond, S. A. et al. Somatodendritic Expression of JAM2 Inhibits Oligodendrocyte Myelination. Neuron 91, 824–836 (2016).

26. Bø, L., Vedeler, C. A., Nyland, H. I., Trapp, B. D. & Mørk, S. J. Subpial demyelination in the cerebral cortex of multiple sclerosis patients. J. Neuropathol. Exp. Neurol. (2003) doi:10.1093/jnen/62.7.723.

27. Howell, O. W. et al. Meningeal inflammation is widespread and linked to cortical pathology in multiple sclerosis. Brain 134, 2755–2771 (2011).

28. Chang, A. et al. Cortical remyelination: A new target for repair therapies in multiple sclerosis. Ann. Neurol. 72, 918–926 (2012).

29. Kidd, D. et al. Cortical lesions in multiple sclerosis. Brain (1999) doi:10.1093/brain/122.1.17.

30. Roosendaal, S. et al. Accumulation of cortical lesions in MS: relation with cognitive impairment. Mult. Scler. J. 15, 708–714 (2009).

31. Nielsen, A. S. et al. Contribution of cortical lesion subtypes at 7T MRI to physical and cognitive performance in MS. Neurology 81, 641–9 (2013).

32. Sacco, R. et al. Cognitive impairment and memory disorders in relapsing–remitting multiple sclerosis: the role of white matter, gray matter and hippocampus. J. Neurol. 262, 1691–1697 (2015).

33. Muhlert, N. et al. The grey matter correlates of impaired decision-making in multiple sclerosis. J. Neurol. Neurosurg. Psychiatry 86, 530–536 (2015).

34. Zonouzi, M. et al. Individual Oligodendrocytes Show Bias for Inhibitory Axons in the Neocortex Individual Oligodendrocytes Show Bias for Inhibitory Axons in the Neocortex. CellReports 27, 2799-2808.e3 (2019).

35. Stedehouder, J. & Kushner, S. A. Myelination of parvalbumin interneurons: A parsimonious locus of pathophysiological convergence in schizophrenia. Mol. Psychiatry 22, 4–12 (2017).

36. Pajevic, S., Basser, P. J. & Fields, R. D. Role of myelin plasticity in oscillations and synchrony of neuronal activity. Neuroscience 276, 135–147 (2014).

37. Orthmann-Murphy, J. et al. Remyelination alters the pattern of myelin in the cerebral cortex. Elife 9, (2020).

38. Olsen, S. R., Bortone, D. S., Adesnik, H. & Scanziani, M. Gain control by layer six in cortical circuits of vision. Nature 483, 47–54 (2012).

39. Denman, D. J. & Contreras, D. Complex Effects on In Vivo Visual Responses by Specific Projections from Mouse Cortical Layer 6 to Dorsal Lateral Geniculate Nucleus. J. Neurosci. 35, 9265–9280 (2015).

40. Bortone, D. S., Olsen, S. R. & Scanziani, M. Translaminar inhibitory cells recruited by layer 6 corticothalamic neurons suppress visual cortex. Neuron 82, 474–485 (2014).

41. Viswanathan, S., Sheikh, A., Looger, L. L. & Kanold, P. O. Molecularly Defined Subplate Neurons Project Both to Thalamocortical Recipient Layers and Thalamus. Cereb. Cortex (2017) doi:10.1093/cercor/bhw271.

42. Clancy, B. & Cauller, L. J. Widespread projections from subgriseal neurons (layer VII) to layer I in adult rat cortex. J. Comp. Neurol. 407, 275–286 (1999).

43. Ibrahim, L. A. et al. Cross-Modality Sharpening of Visual Cortical Processing through Layer-1-Mediated Inhibition and Disinhibition. Neuron 89, 1031–1045 (2016).

44. Madisen, L. et al. A robust and high-throughput Cre reporting and characterization system for the whole mouse brain. Nat. Neurosci. (2010) doi:10.1038/nn.2467.

45. Gong, S. et al. Targeting Cre recombinase to specific neuron populations with bacterial artificial chromosome constructs. Journal of Neuroscience vol. 27 9817–9823 (2007).

46. Harris, J. A. et al. Anatomical characterization of Cre driver mice for neural circuit mapping and manipulation. Front. Neural Circuits (2014) doi:10.3389/fncir.2014.00076.

47. Zonouzi, M. et al. Individual Oligodendrocytes Show Bias for Inhibitory Axons in the Neocortex. Cell Rep. 27, 2799-2808.e3 (2019).

48. Hunt, B. A. E. et al. Relationships between cortical myeloarchitecture and electrophysiological networks. Proc. Natl. Acad. Sci. 113, 13510–13515 (2016).

49. Nieuwenhuys, R. & Broere, C. A. J. A detailed comparison of the cytoarchitectonic and myeloarchitectonic maps of the human neocortex produced by the Vogt–Vogt school. Brain Struct. Funct. 1, 3 (2020).

50. Whissell, P. D., Cajanding, J. D., Fogel, N. & Kim, J. C. Comparative density of CCK- and PV-GABA cells within the cortex and hippocampus. Front. Neuroanat. 9, (2015).

51. Mayoral, S. R., Etxeberria, A., Shen, Y. A. A. & Chan, J. R. Initiation of CNS Myelination in the Optic Nerve Is Dependent on Axon Caliber. Cell Rep. (2018) doi:10.1016/j.celrep.2018.09.052.

52. Stedehouder, J. et al. Local axonal morphology guides the topography of interneuron myelination in mouse and human neocortex. Elife 8, 1–28 (2019).

53. Snaidero, N. et al. Myelin replacement triggered by single-cell demyelination in mouse cortex. Nat. Commun. 11, 4901 (2020).

54. Bacmeister, C. M. et al. Motor learning promotes remyelination via new and surviving oligodendrocytes. Nat. Neurosci. 23, 819–831 (2020).

55. Schain, A. J., Hill, R. A. & Grutzendler, J. Label-free in vivo imaging of myelinated axons in health and disease with spectral confocal reflectance microscopy. Nat. Med. 20, 443– 449 (2014).

56. Lee, S. et al. A culture system to study oligodendrocyte myelination processes using engineered nanofibers. Nat. Methods 9, 917–922 (2012).

57. Kang, S. H., Fukaya, M., Jason, Y. K., Rothstein, J. D. & Bergles, D. E. NG2+ CNS glial progenitors remain comitted to the oligodendrocyte lineage in postnatal life and following neurodegenration. Neuron 68, 668–81 (2010).

58. Psachoulia, K., Jamen, F., Young, K. M. & Richardson, W. D. Cell cycle dynamics of NG2 cells in the postnatal and ageing brain. Neuron Glia Biol. 5, 57 (2009).

59. Young, K. M. et al. Oligodendrocyte dynamics in the healthy adult CNS: Evidence for myelin remodeling. Neuron 77, 873–885 (2013).

60. Kang, S. H., Fukaya, M., Yang, J. K., Rothstein, J. D. & Bergles, D. E. NG2+ CNS glial progenitors remain committed to the oligodendrocyte lineage in postnatal life and following neurodegeneration. Neuron 68, 668–681 (2010).

61. Zhu, X. et al. Age-dependent fate and lineage restriction of single NG2 cells. Development 138, 745–753 (2011).

62. Welliver, R. R. et al. Muscarinic Receptor M <sub>3</sub>R Signaling Prevents Efficient Remyelination by Human and Mouse Oligodendrocyte Progenitor Cells. J. Neurosci. 38, 6921–6932 (2018).

63. Zuchero, J. B. et al. CNS Myelin Wrapping Is Driven by Actin Disassembly. Dev. Cell 34, 152–167 (2015).

64. Snaidero, N. et al. Myelin membrane wrapping of CNS axons by PI(3,4,5)P3-dependent polarized growth at the inner tongue. Cell 156, 277–290 (2014).

65. Ishibashi, T. et al. Astrocytes promote myelination in response to electrical impulses. Neuron 49, 823–832 (2006).

66. Mitew, S. et al. Pharmacogenetic stimulation of neuronal activity increases myelination in an axon-specific manner. Nat. Commun. 9, 1–16 (2018).

67. Hines, J. H., Ravanelli, A. M., Schwindt, R., Scott, E. K. & Appel, B. Neuronal activity biases axon selection for myelination in vivo. Nat. Neurosci. 18, 683–689 (2015).

68. Xin, W. & Chan, J. R. Myelin plasticity: sculpting circuits in learning and memory. Nature Reviews Neuroscience vol. 21 682–694 (2020).

69. Dutta, D. J. et al. Regulation of myelin structure and conduction velocity by perinodal astrocytes. Proc. Natl. Acad. Sci. U. S. A. 115, 11832–11837 (2018).

70. Hughes, E. G., Kang, S. H., Fukaya, M. & Bergles, D. E. Oligodendrocyte progenitors balance growth with self-repulsion to achieve homeostasis in the adult brain. Nat. Neurosci. 16, 668–676 (2013).

71. Ghanbari, L. et al. Cortex-wide neural interfacing via transparent polymer skulls. doi:10.1038/s41467-019-09488-0.

72. Hamada, M. S. & Kole, M. H. P. Myelin Loss and Axonal Ion Channel Adaptations Associated with Gray Matter Neuronal Hyperexcitability. J. Neurosci. 35, 7272–7286 (2015).

73. Hamada, M. S., Popovic, M. A. & Kole, M. H. P. Loss of Saltation and Presynaptic Action Potential Failure in Demyelinated Axons. Front. Cell. Neurosci. 11, 1–11 (2017).

74. Werneburg, S. et al. Targeted Complement Inhibition at Synapses Prevents Microglial Synaptic Engulfment and Synapse Loss in Demyelinating Disease. Immunity 52, 167-182.e7 (2020).

75. Rankin, K. A. et al. Selective estrogen receptor modulators enhance CNS remyelination independent of estrogen receptors. J. Neurosci. 39, 2184–2194 (2019).

76. Mei, F. et al. Accelerated remyelination during inflammatory demyelination prevents axonal loss and improves functional recovery. Elife 5, (2016).

77. Mei, F. et al. Identification of the Kappa-Opioid Receptor as a Therapeutic Target for Oligodendrocyte Remyelination. (2016) doi:10.1523/JNEUROSCI.1493-16.2016.

78. Taniguchi, H. et al. A Resource of Cre Driver Lines for Genetic Targeting of GABAergic Neurons in Cerebral Cortex. Neuron 71, 995–1013 (2011).

79. Longair, M. H., Baker, D. A. & Armstrong, J. D. Simple Neurite Tracer: open source software for reconstruction, visualization and analysis of neuronal processes. Bioinformatics 27, 2453–4 (2011).

80. Hoffmann, H. violin.m -Simple violin plot using matlab default kernal density estimation. (2015).

81. Papadopoulos, G. C., Parnavelas, J. G. & Buijs, R. M. Light and electron microscopic immunocytochemical analysis of the serotonin innervation of the rat visual cortex. J. Neurocytol. (1987) doi:10.1007/BF01611992.

82. Papadopoulos, G. C., Parnavelas, J. G. & Buijs, R. M. Light and electron microscopic immunocytochemical analysis of the noradrenaline innervation of the rat visual cortex. J. Neurocytol. 18, 1–10 (1989).

83. Papadopoulos, G. C., Parnavelas, J. G. & Buijs, R. M. Light and electron microscopic immunocytochemical analysis of the dopamine innervation of the rat visual cortex. J. Neurocytol. 18, 303–310 (1989).

